# microRNAs control temporal patterning of the vertebrate nervous system

**DOI:** 10.64898/2026.10.02.755877

**Authors:** Estelle Renaux, Laia Caudet Segarra, Archi Shailesh Mistry, Manish Pandey, Julia Sagner, Hans-Martin Jäck, Jürgen Wittmann, Michael Wegner, Andreas Sagner

**Affiliations:** Institut für Biochemie, Friedrich-Alexander-Universität Erlangen-Nürnberg (FAU), Erlangen, Germany; Division of Molecular Immunology, Department of Internal Medicine III, Nikolaus-Fiebiger-Center for Molecular Medicine (NFZ), Friedrich-Alexander-Universität Erlangen-Nürnberg (FAU), Erlangen, Germany; Department of Translational Immunology, Department of Medicine 3 – Rheumatology and Immunology, Nikolaus-Fiebiger-Center for Molecular Medicine (NFZ), Friedrich-Alexander-Universität Erlangen-Nürnberg (FAU), Erlangen, Germany; Deutsches Zentrum Immuntherapie, Friedrich-Alexander-Universität Erlangen-Nürnberg (FAU) and Uniklinikum Erlangen, Erlangen, Germany

## Abstract

Temporal patterning is essential for generation of neuronal diversity in the developing vertebrate nervous system. A shared sequence of temporal transcription factors (tTFs) partitions neurons by birth date across much of the central nervous system. The molecular mechanisms that control the timing of expression of these tTFs are incompletely understood. Here, we provide evidence that this process is under the control of microRNAs (miRNAs). Perturbing miRNA biogenesis by conditional deletion of Dgcr8 from the developing mouse nervous system extends the window during which neurons with early identities are generated, but abolishes late identities in the spinal cord, hindbrain and midbrain, and holds progenitors in an early transcriptional state. We further identify miRNAs of the let-7 and miR-9 families, which target transcriptional regulators of early progenitor and neuronal identities, as candidates for mediating this effect. Throughout the nervous system, levels of these miRNAs rise during the neurogenic period in progenitors, while levels in neurons correlate with their birth-date. Moreover, the sensitivity of tTF-encoding transcripts to miR-9 inversely correlates with their expression sequence, suggesting a model in which miR-9 controls the chronology of neuronal cell fate transitions by targeting specifiers of early progenitor and neuronal temporal identity in a level-dependent manner. Consistently, perturbations of miR-9 expression dynamics by gain and loss-of-function approaches perturb temporal patterning.

Together, our results demonstrate a critical, evolutionary conserved role for miRNAs in temporal patterning and suggest that a temporal miR-9 expression gradient controls the timing of tTF expression in large parts of the developing vertebrate nervous system.

## Introduction

A fundamental question in neuroscience is how the intricate complexity of the central nervous system (CNS) arises during development. Recent advances have uncovered a surprisingly large number of neuronal subtypes, defined by unique molecular profiles and specialized functions. These findings raise a critical issue: what developmental processes and molecular mechanisms underlie the precise generation of this neuronal diversity, ensuring that specific neuronal subtypes are produced at the right time, in the right place, and in the appropriate numbers? Over the last few decades, considerable attention has been given to spatial patterning mechanisms, which subdivide the embryonic nervous system into defined topographical regions. These spatial programs direct the emergence of distinct neuronal classes that share molecular hallmarks and execute similar functions along the dorsal-ventral (DV) and rostral-caudal axes of the neural tube (*1–3*). However, even within a single region of the nervous system, these neuronal classes can typically be further partitioned into distinct molecular and functional subtypes.

The sequential generation of distinct cell types from the same progenitor population contributes to neuronal diversity in several developmental systems, including the insect embryo and visual system and the vertebrate cortex, retina and spinal cord (*4–7*). Temporal patterning of large regions of the developing vertebrate nervous system is mediated at least in part by a series of transcription factors (TFs) that are expressed in a chronological sequence, with each TF marking neurons born at specific developmental windows (*8*, *9*). As neurogenesis progresses, the sequential activation of these TFs assigns temporal identities to neurons, thereby further diversifying the output of spatially defined progenitor domains. In the mouse spinal cord, for instance, neurons generated around embryonic day (e)9.5 express members of the *Onecut* (*OC*) TF family. Neurons generated between e9.5 and e11.5 express *Zinc finger homeobox 2-4* (*Zfhx2-4)* and can be further subdivided into *POU-class 2 homeobox 2 (Pou2f2)* expressing early-intermediate neurons, produced around e10.5, and *POU-class 3 homeobox 1 (Pou3f1)* expressing late-intermediate neurons, produced around e11.5 (*10–12*). Finally, late-born neurons, produced between e11.5 and e13.5, express *Nuclear factor I A/B/X* (*Nfia/b/x), B-Cell CLL/Lymphoma 11B (Bcl11b*, also known as *Ctip2)* and *Neuronal differentiation 2/6* (*Neurod2/6)* (*8*, *10*, *12*, *13*).

Growing evidence points to the functional relevance of this temporal program. Members of the OC TF family are essential for generating various neuronal subtypes throughout the CNS, including the spinal cord (*14–19*), while Pou2f2 and Pou3f1 are essential for the correct generation of anterolateral system and motor neuron subtypes during spinal cord development (*11*). Expression of *Zfhx3/4* or *Nfib/Neurod2* moreover correlates with long-range projection or local interneuron identities, respectively (*12*), and Neurod2/6 control neuropeptide expression in late-born spinal inhibitory dorsal horn neurons (*20*). Similar roles for these TFs have been observed in other parts of the CNS such as the retina, where OC, Pou2f2, and Nfi family members mark early- and late-born neuronal subtypes and are required for their development (*16*, *21–23*). These findings indicate that this conserved temporal transcription factor (tTF) code plays a critical role in specifying neuronal subtypes across extensive regions of the nervous system. However, the molecular mechanisms controlling this temporal program are still poorly understood.

MicroRNAs (miRNAs) are short RNA molecules, averaging 22 nucleotides in length, that regulate gene expression by binding to the 3′ untranslated regions (3’-UTRs) of target mRNAs, thereby promoting mRNA degradation and inhibiting translation (*24–27*). Most miRNAs are transcribed by RNA polymerase II or III as primary transcripts (pri-miRNAs). In the canonical and predominant miRNA biogenesis pathway, pri-miRNAs are processed in the nucleus by the microprocessor complex into precursor miRNAs (pre-miRNAs). This complex comprises RNA-binding protein DiGeorge Syndrome Critical Region 8 (Dgcr8), which recognizes specific sequence motifs within the pri-miRNA, and RNase III enzyme Drosha, which cleaves the miRNA hairpin structure at its base to generate the pre-miRNA (*28–33*). Pre-miRNAs are then exported to the cytoplasm, where they are further processed by the Dicer-TRBP complex, which removes the terminal loop to generate a miRNA duplex (*34*, *35*). This duplex is subsequently loaded onto Argonaute proteins to form the RNA-induced silencing complex (RISC) (*36–38*), which unwinds the miRNA duplex into single-stranded miRNAs. One strand, retained by RISC, becomes the functional mature miRNA, while the opposite strand is degraded. The directionality of the retained strand determines whether the mature miRNA is designated as the 5p or 3p form. For any given miRNA, the relative abundance of 5p versus 3p strands can vary widely across cell types and physiological conditions, ranging from approximately equal proportions to a strong dominance of one strand (*25*, *26*).

miRNAs are implicated in temporal patterning of different brain regions. In the mammalian neocortex, miR-128, miR-9, and let-7 establish opposing, functionally antagonistic temporal expression and activity gradients, which together regulate cortical layer formation (*39*, *40*). Similarly, let-7, miR-125, and miR-9, are key regulators of the early-to-late developmental transition in retinal progenitors (*41*, *42*). In *Drosophila* miRNAs transcribed from the *let-7-C* locus regulate temporal cell fate in the mushroom body lineage (*43*). In the spinal cord, miRNAs are required for the switch from neuro-to gliogenesis and have been implicated in the generation of distinct motor neuron subtypes (*44–47*). These observations led us to ask whether miRNAs provide a general timing mechanism for temporal patterning across the developing CNS.

Here, we show that inhibition of miRNA biogenesis by *Dgcr8* deletion disrupts progenitor maturation and causes an increase in early-born neuronal populations and near-complete absence of neurons with late-born identities in the mouse embryonic midbrain, hindbrain and spinal cord. Computational analysis identified miRNAs of the miR-9 and let-7 families as regulators mediating this effect by evolutionary conserved targeting of several TFs implicated in early progenitor development and early neuronal tTFs. In embryos and stem cell-derived neuronal differentiation paradigms, miR-9 and let-7 levels gradually increase over the neurogenic period, while the sensitivity of neuronal tTF-encoding transcripts to miR-9 inversely correlates with their order of expression during development. Consistently, perturbing miR-9 expression dynamics by gain- and loss-of-function studies in chicken embryos disrupts temporal patterning. Together, our results demonstrate a critical and evolutionary conserved role for miRNAs in temporal patterning and suggest that a temporal miR-9 expression gradient controls the timing of tTF expression in large parts of the developing vertebrate nervous system.

## Results

### Perturbation of miRNAs biogenesis disrupts temporal patterning of the CNS

To assess whether miRNAs are required for temporal patterning, we perturbed miRNA biogenesis by conditional deletion of *Dgcr8* using the *Brn4-Cre* (also known as *Tg(Pou3f4)-Cre*) line (*48*, *49*), which recombines in large parts of the developing nervous system before the onset of neurogenesis. Cre expression in the spinal cord of *Brn4-Cre; Dgcr8 flox/flox* embryos (hereafter called Dgcr8 cKO embryos) initiated around e9.5, Dgcr8 was undetectable in neurons and progenitors by e11.5 (Fig. S1A). To assess the consequences on temporal neuronal subtype specification, we stained e13.5 spinal cord sections for the intermediate neuron markers Pou2f2 and Zfhx3 and the late neuronal markers Neurod2 and Bcl11b. Although the total number of nuclei detected in Dgcr8 cKO sections was significantly reduced, Pou2f2 and Zfhx3-positive neurons were significantly increased and the late neuronal markers Bcl11b and Neurod2 essentially absent in Dgcr8 cKO embryos (Fig.1A-B). The fraction of Zfhx3-positive neurons that co-expressed the early intermediate marker Pou2f2 also increased, suggesting that the temporal program may not advance beyond the early-intermediate Pou2f2 stage in these mutants (Fig. 1A-C). Consistent with this, a more fine-grained analysis using OC2, Pou2f2 and Pou3f1 confirmed an increase in neurons expressing the early and early-intermediate tTFs OC2 and Pou2f2. In contrast, the number of neurons single-positive for the late-intermediate tTF Pou3f1 decreased, with many Pou3f1-positive neurons co-expressing either OC2, Pou2f2 or both markers (Fig. 1D-E). Taken together, these results suggest an essential requirement for miRNAs to advance the temporal program beyond the earliest temporal identities in the embryonic spinal cord.

**Fig. 1.**
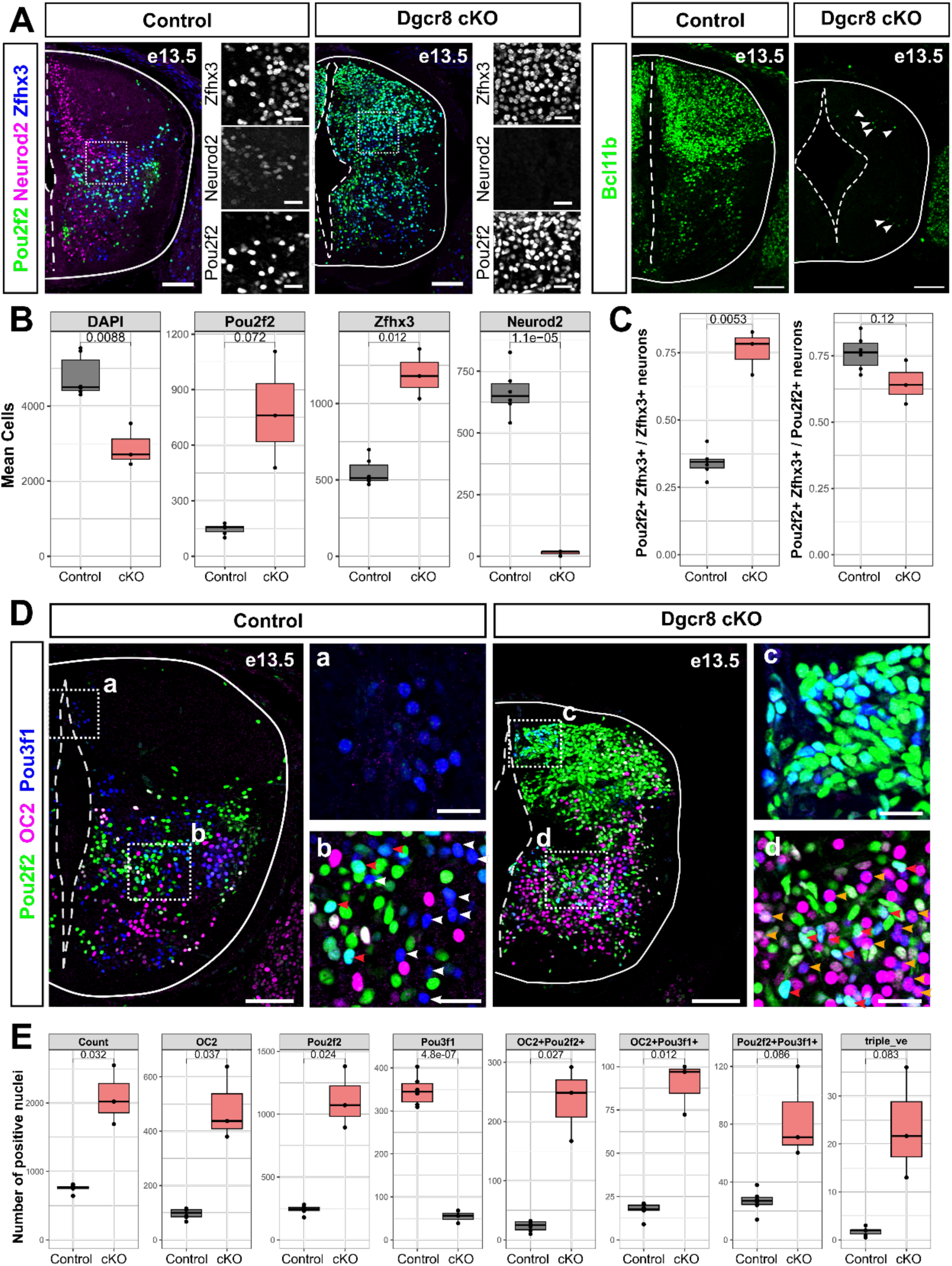
Dgcr8 deletion disrupts temporal patterning of the vertebrate spinal cord. **(A)** E13.5 cervical spinal cord sections of control (left) and Dgcr8 cKO embryos (right) stained for Pou2f2, Zfhx3, Neurod2 and Ctip2. **(B)** Average number of cells per spinal cord section stained for the respective markers. **(C)** Fraction of all Zfhx3-positive (left) and Pou2f2-positive (right) neurons that co-express both markers. **(D)** E13.5 cervical spinal cord sections of control (left) and Dgcr8 cKO embryos (right) stained for OC2, Pou2f2, and Pou3f1. White arrowheads point to single-positive Pou3f1^+^ neurons, red arrowheads point to Pou2f2^+^Pou3f1^+^ double-positive neurons, and orange arrowheads point to OC2^+^Pou3f1^+^ double-positive neurons. **(E)** Quantification of the number of nuclei positive for OC2, Pou2f2, and Pou3f1 TF combinations. Scale bars = 100 μm, insets = 25 μm, Statistical significance in B, C and E was assessed by Student’s t-test. p-values are integrated on the graphs.

We next investigated whether loss of Dgcr8 affects temporal patterning in other CNS regions. Similar to the spinal cord, Dgcr8 recombination was nearly complete in the midbrain by e11.5 and loss of Dgcr8 caused an increase in early OC2 neurons and decrease of Pou3f1 neurons. This effect was even more pronounced at e13.5 (Fig. S1B-C). However, in contrast to the spinal cord, the amount of Pou2f2 and Zfhx3 neurons was also markedly reduced in the midbrain, suggesting that the loss of Dgcr8 affects temporal progression in the midbrain more severely than in the spinal cord (Fig. S1C-D). Moreover, the overall structure of the midbrain was severely disrupted in the Dgcr8 cKO embryos, with pronounced decrease in midbrain size that became particularly evident at e13.5 (Fig. S1B-D), suggesting that besides alterations to temporal patterning, other effects contribute to the observed phenotype. The phenotype in the hindbrain largely resembled our observations in the spinal cord. Loss of Dgcr8 resulted in an expansion of early OC2 and early-intermediate Pou2f2 and Zfhx3 populations, whereas the number of neurons expressing the late-intermediate and late markers Pou3f1 and Nfib were markedly reduced (Fig. S1E-F). Collectively, these findings identify a critical role for miRNAs in temporal patterning of the midbrain, hindbrain and spinal cord. In combination with previous observations in the retina and forebrain (*39–41*, *50*), these observations suggest that miRNAs are critical for temporal patterning throughout the developing CNS.

### Characterization of growth and neuronal differentiation dynamics of the embryonic Dgcr8 cKO spinal cord

To determine whether the observed differences in early and late-born neuronal subtype numbers are a consequence of altered growth or neuronal differentiation dynamics, we focused on the embryonic spinal cord and quantified the size of different spinal cord compartments (total area, neuronal area, progenitor area, and central canal area), apoptosis and proliferation. At e11.5, the size of the different compartments did not differ between control and mutant embryos (Fig. S2A, D). Similarly, co-detection of the progenitor marker Sox2 and the late neurogenic/neuronal marker Sox11 (*51*) revealed no significant changes in the numbers of Sox2⁺ and Sox2⁺ Sox11⁺ cells, indicating that the number of progenitors and neurogenic cells at this stage is unaltered (Fig. S2E–F). In contrast, at e13.5, total spinal cord area and neuronal area were significantly reduced in the mutants, while the central canal area was significantly increased (Fig. S2G-H). The ventricular zone showed a slight, although non-significant increase in area, but often appeared discontinuous and did not line the entire lumen of the central canal (Fig. S2G-H). Cleaved caspase 3 (cC3) staining revealed a significant increase in apoptotic cells in Dgcr8 cKO embryos compared to controls in the ventricular and marginal zones at e11.5 and e13.5 (Fig. S2A-C, I-J). Mitotic activity, assessed by phospho-histone H3 (pHH3) immunostaining, was not altered at e11.5, but significantly increased at e13.5 (Fig. S2A, B, I, K). Collectively, these data suggest that the loss of miRNAs causes changes in tissue morphology, proliferative behavior and increased apoptosis. However, the overall effect of these changes was largely restricted to the late neurogenic phase, suggesting the strong reduction of late temporal identity neurons and increase in early identity neurons in the Dgcr8 cKO embryos is not due to altered temporal neurogenesis patterns or precocious depletion of neural progenitors. The observed effects on tissue morphology and apoptosis are therefore unlikely to fully account for the temporal patterning phenotype of the Dgcr8 cKO embryos.

We next tested whether loss of miRNAs differentially affects survival of neurons born at different stages during embryonic development. To this end, we performed neuronal birth-dating by administering 5’-ethynyl-2′-deoxyuridine (EdU) at e10.5 or e11.5 and quantified the number of EdU+ nuclei at e13.5. Spinal cord sections from control and Dgcr8 cKO embryos treated with EdU either at e10.5 or e11.5 had comparable nuclei numbers to un-injected embryos of the same genotype, suggesting EdU-administration does not impact cell survival (Fig. 2A, B, Fig 1B). In control embryos, roughly similar numbers of neurons were labelled by EdU administered at e10.5 or e11.5 (Fig. 2B, E). In contrast, the total number of EdU+ nuclei in Dgcr8 cKO embryos was slightly reduced when EdU was administered at e10.5, but strongly reduced when EdU was given at e11.5 (Fig. 2A, B, D, E). Because overall (Sox2+) and neurogenic (Sox2+ Sox11+) progenitor numbers were unaltered at e11.5 (Fig. S2E–F), these findings suggest that the survival of neurons born after e10.5 is severely diminished in the Dgcr8 cKO embryos.

**Fig. 2.**
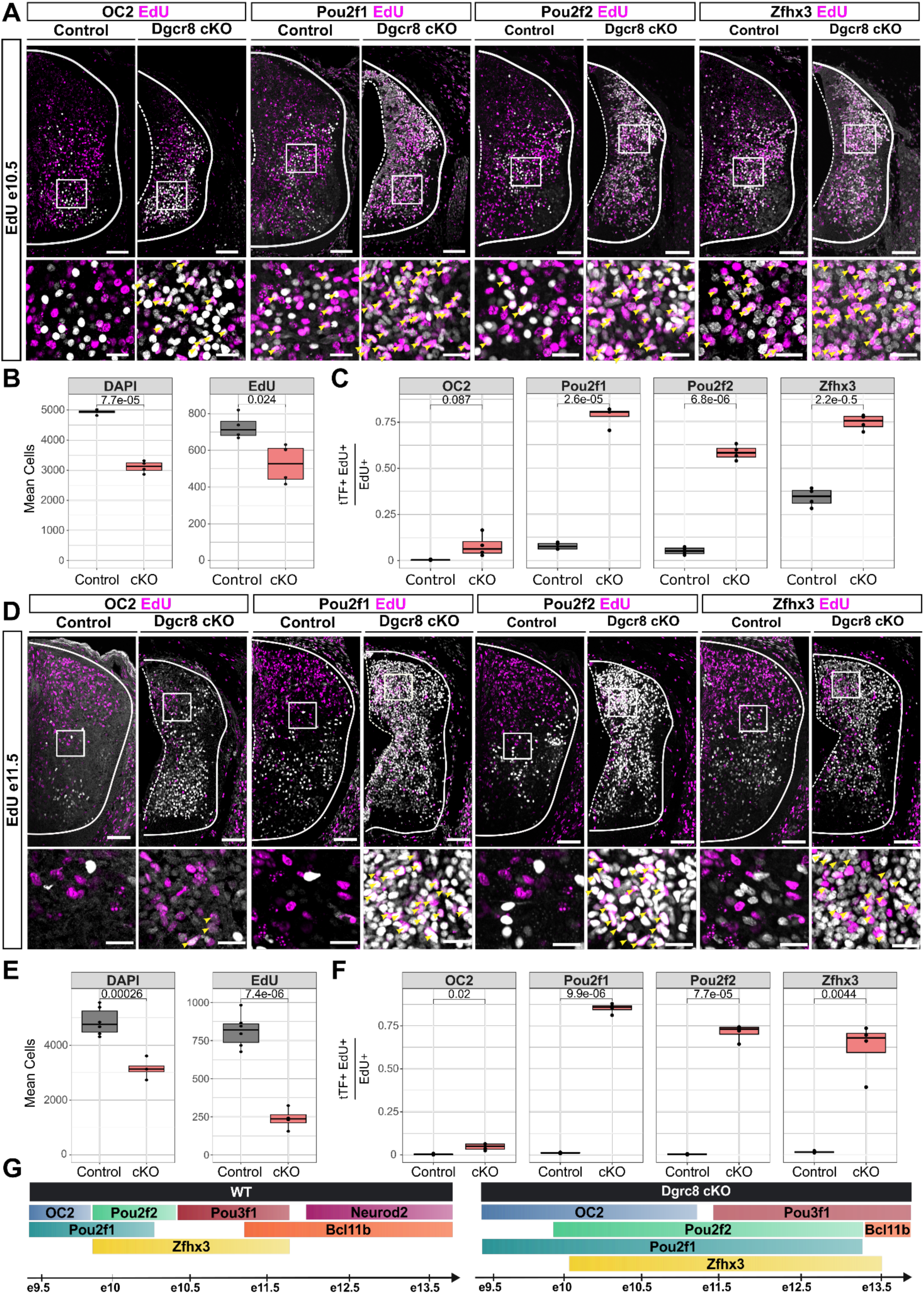
EdU birth-dating confirms chronological shifts in the generation of early tTF expressing neurons. **(A-C)** Analysis of e13.5 spinal cord sections, when EdU was administered at e10.5**. (A)** Sections of control and Dgcr8 cKO embryos co-stained for OC2, Pou2f1, Pou2f2, and Zfhx3 and EdU. **(B)** Quantification of the total number (DAPI) and EdU+ nuclei. **(C)** Fraction of EdU+ neurons that express OC2, Pou2f1, Pou2f2 and Zfhx3. **(D-F)** Analysis of e13.5 spinal cord sections, when EdU was administered at e11.5**. (D)** Sections of control and Dgcr8 cKO embryos co-stained for OC2, Pou2f1, Pou2f2, and Zfhx3 and EdU. **(E)** Quantification of the total number (DAPI) and EdU+ nuclei. **(F)** Fraction of EdU+ neurons that express OC2, Pou2f1, Pou2f2 and Zfhx3. **(G)** Schematic summarizing the observed changes in tTF expression between control (left) and Dgcr8 cKO embryos (right). Scale bars = 100 μm, insets = 25 μm, Statistical significance in B, C, E, F was assessed by Student’s t-test. p-values are integrated on the graphs.

While the reduced survival provides an explanation for the reduction of late-born neurons, it does not explain the increase in neurons expressing early tTFs in the Dgcr8 cKO embryos. We therefore hypothesized that loss of miRNAs extends the developmental window, during which neurons expressing these early tTFs are generated. To test this, we quantified the fraction of EdU+ nuclei labelled by the early tTFs OC2, Pou2f1, Pou2f2, and Zfhx3. Consistent with previous observations (*8*, *11*), only a minor fraction of EdU+ nuclei expressed OC2, Pou2f1 or Pou2f2, which are specific for neurons born before e10.5, in control embryos, when EdU was injected at e10.5 or e11.5 (Fig. 2A, C. D, F). Zfhx3 was present in 30-40% of EdU+ nuclei, when EdU was administered at e10.5; however, this fraction approached 0, when EdU was injected at e11.5 (Fig. 2A, C, D, F). In the Dgcr8 cKO embryos, the fraction of EdU+ nuclei labelled by each of these early tTFs increased (Fig. 2A, C, D, F). While the increase for OC2 was small, the fraction of EdU+ nuclei labelled by Pou2f1, Pou2f2 and Zfhx3 increased for both injection timepoints to 60-80% (Fig. 2A, C, D, F). Pou2f1 expression appeared particularly expanded and was unlike Pou2f2 and Zfhx3 detected in the ventricular zone, suggesting miRNAs are directly or indirectly repressing Pou2f1 in progenitors (Fig. 2A, D). These observations provide experimental evidence for an expansion of the developmental windows during which early tTF-expressing neurons are generated in the Dgcr8 cKO embryos (Fig. 2G). They also raise the possibility that loss of miRNAs not only perturbs the temporal identity of neurons, but also affects transcriptional programs in neural progenitors.

### Characterization of temporal patterning defects in Dgcr8 cKO embryos by scRNAseq

To characterize the transcriptional changes caused by loss of miRNA biosynthesis and gain insights into the underlying molecular mechanisms, we performed single-cell RNA sequencing (scRNAseq) analysis on Dgcr8 cKO and control spinal cords at e11.25 (3 controls, 3 mutants, 26019 cells) and e13.25 (3 controls, 4 mutants, 39971 cells) (Fig. 3A). After quality control, filtering and computational doublet removal, datasets were integrated using the Seurat v5 CCA workflow and clustered based on transcriptional similarity (see Supplemental Experimental Procedures). Differential gene expression (DGE) and marker-based analyses enabled the identification of major spinal cord cell populations, including dorsal and ventral neuronal classes, progenitor populations, and cell types not derived from the spinal cord, which were removed from subsequent analyses. Iterative sub-clustering further refined heterogeneous populations and improved segregation of closely related neuronal and progenitor identities. These curated annotations were subsequently used to compare transcriptional and cellular changes between control and Dgcr8 cKO embryos (Fig. 3A). Overall, marker gene expression remained well-segregated between the cardinal classes of neurons at e11.25 and e13.25 (Fig. S3A,B), however, at e13.25 the relative fraction of neurons in these classes was altered in the Dgcr8 cKO mutants (Fig. S3C). Consistent with the observed reduction of late-born neurons (Fig. 1), the contribution of neurons to the predominantly late-born inhibitory dI4/dILA and excitatory dI5/dILB classes that form the dorsal horns of the spinal cord was reduced, while other neuronal populations were unchanged or even slightly increased (Fig. S3C). Together, these results suggest that miRNA biogenesis is dispensable for the establishment of broad spinal cord cell-type identities.

**Fig. 3.**
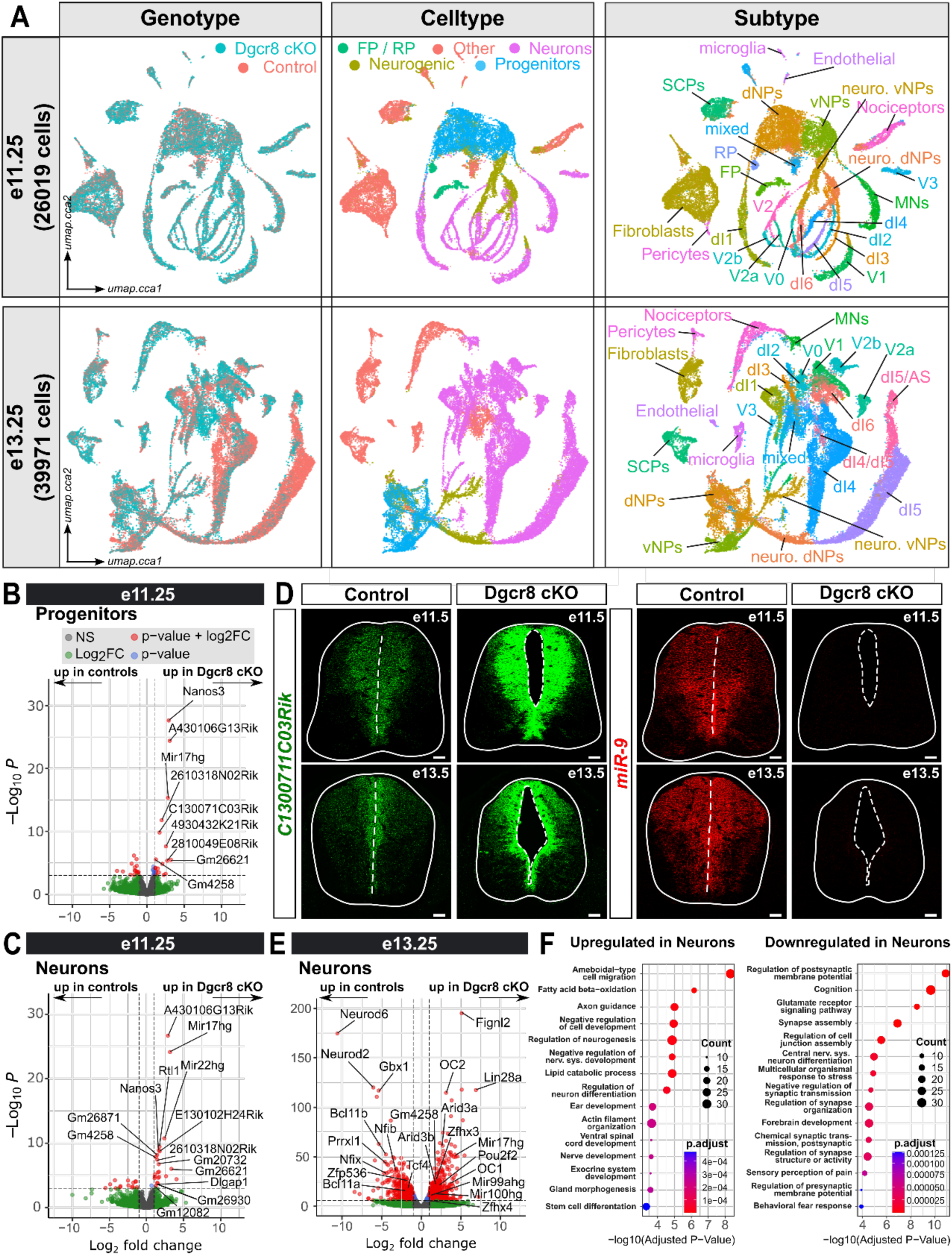
Characterization of temporal patterning defects in Dgcr8 cKO embryos by scRNAseq. **(A)** UMAP representation of scRNAseq data from e11.25 (top row) and e13.25 (bottom row) embryos color-coded for genotype (left), broad cell type (middle) and specific subtype identities. **(B-C)** DGE analysis comparing gene expression in e11.25 neural progenitors (B) and neurons (C) between control and Dgcr8 cKO embryos. **(D)** miR-9 (red, left) is absent in e11.5 (top row) and e13.5 (bottom row) spinal cord sections from Dgcr8 cKO embryos, while levels of the miR-9 host transcript *C130071C03Rik* (green, right) are elevated in these embryos. **(E)** DGE analysis of e13.25 neurons. **(F)** GO terms associated with up-(left) and down-regulated transcripts in the Dgcr8 cKO embryos. Scale bars in D = 100 µm

To understand how the temporal patterning phenotype of the Dgcr8 cKO embryos emerges, we first focused on the e11.25 datasets and performed DGE analysis using a DESeq2-based pseudobulk approach to compare gene expression in neurons and progenitors between Dgcr8 cKO and control embryos. The most prominent effect was an upregulation of pri-miRNAs in neurons and progenitors in the Dgcr8 cKO animals. These included the miR-181 family pri-miRNAs *Nanos3*, *A430106G13Rik* and *Gm4258*, miR-9 family pri-miRNA *C130071C03Rik*, and pri-miRNAs derived from the *Mir17hg* miRNA cluster and the let-7 miRNA containing clusters *Mir99ahg* and *Mir100hg* (Fig. 3B-C). To experimentally validate this observation, we performed RNAscope for miR-9 and its pri-miRNA transcript *C130071C03Rik* on e11.5 and e13.5 spinal cord sections. Consistent with the predictions of the DGE analysis, *C130071C03Rik* levels were upregulated in the Dgcr8 cKO embryos, while miR-9 was nearly undetectable (Fig. 3D). We next focused on the expression of TFs at e11.25. Pseudobulk DGE analysis did not reveal any de-regulated TF transcripts. Similarly, inspecting the expression dynamics of the neuronal tTFs did not reveal any striking differences (Fig. S3D). Collectively, these data suggest that consistent with the described role of Dgcr8 in the miRNA biogenesis pathway, pri-miRNA transcripts are upregulated and mature miRNAs reduced in Dgcr8 cKO embryos at e11.25, but no major temporal patterning phenotypes can be detected at the transcriptional level.

We next focussed on neurons at e13.25. DGE analysis revealed marked shifts in neuronal temporal identity in the Dgcr8 cKO mutants. Early temporal identity markers such as *OC1aaaOC2*, *Pou2f2*, *Zfhx3,* and *Lin28a* were upregulated, while late temporal markers such as *Bcl11a, Bcl11b*, *Neurod2, Neurod6, Tcf4* and *Nfib,* and dorsal horn markers such as *Sall3, Gbx1, Lmx1b, Tlx3* and *Prrxl1* were downregulated (Fig. 3E, Fig. S4A-B). To gain a more-systematic overview over the biological processes perturbed by miRNA loss, we performed Gene Ontology (GO) analysis. Genes upregulated in Dgcr8 cKO neurons were associated with GO terms such as axon guidance, neurogenesis, neuron differentiation, actin filament organization, and ventral spinal cord development, typically associated with earlier stages of neuronal circuit formation. In contrast, downregulated genes were enriched for terms related to synapse assembly and function, cognition, and sensory perception of pain (Fig. 3F). Collectively, these data confirm the previously observed changes in temporal patterning and the numbers of neurons in different neuronal classes and suggest a requirement for miRNAs for neuronal maturation and circuit formation of spinal cord neurons.

### Loss of Dgcr8 affects chronological neuronal subtype specification within cardinal classes

Chronological neuronal subtype generation has been previously described in different cardinal classes of spinal cord neurons (*10*, *12*). Examples include the sequential generation of excitatory dI5 and dILB neurons, which largely correlates with their subdivision into early-born projection neurons of the anterolateral system (AS), marked by embryonic expression of the TF Phox2a, and later-born local interneurons (*13*, *52*, *53*). Similarly, sequential generation of different clades of inhibitory V1 and excitatory V3 neurons has been described, with these populations contributing to the control of different aspects of coordinated locomotion (*14*, *54*– *56*). How temporal patterning of these neuronal classes is controlled and whether it is under the control of the same upstream regulators, including miRNAs, as the global tTFs is still largely unclear.

We therefore asked whether loss of Dgcr8 affects temporal identity specification of these neuronal classes. Within the dI5/dILB population, markers of early-born long-range projection AS neurons, including *Phox2a* and *Zfhx3*, were increased in Dgcr8 cKO embryos, whereas markers associated with later-born local projection neurons, such as *Prrxl1, Rora* and *Tcf4*, were decreased (Fig. S4C). In V1 neurons, markers for the early-born Renshaw and Pou6f2 subtypes, including *OC1*, *OC2*, *MafA*, *MafB*, *Calb1*, *Pou2f2* and *Pou6f2,* were upregulated, whereas markers for the later-born Foxp2 and Sp8 clades, including *Foxp2*, *Zfpm2*, *Esrrg*, *Ebf1*, *Nfib* and *Neurod2* were downregulated (Fig. S4A, B, D) (*14*, *57*). In V3 neurons, markers associated with early-born dorsal V3 (V3d) neurons, including *Lhx1*, were increased, whereas markers characteristic of later-born ventral V3 (V3v) neurons, such as *Olig3*, were reduced in the mutants (Fig. S4E) (*11*, *54*).

These findings were confirmed by immunostainings of e13.5 spinal cord sections. Consistent with the scRNAseq results (Fig. S4C-E), the overall number of dI5/dILB neurons, labelled by Lmx1b, was significantly reduced. However, despite this reduction, the number of Phox2a+ and Zfhx3+ dI5/dILB neurons increased in the Dgcr8 cKO embryos, suggesting a large fraction of the remaining dI5/dILB neurons acquired an AS neuron identity (Fig. S4F-G). In V1 neurons, labelled by En1, Pou2f2+ and Calb1+ cells were increased, while Foxp2+ cells were strongly reduced (Fig. S4H-I). Similarly, Pou2f2+ cells were significantly increased in V3 neurons, labelled by Nkx2.2, while Olig3+ V3 neurons were essentially absent. Interestingly, we did not detect a corresponding change in the number of Lhx1-positive V3 neurons, suggesting that Lhx1 protein levels in V3 neurons do not follow global tTF expression and are regulated independently of miRNAs (Fig. S4J-K). Collectively, these findings suggest that miRNAs are required for the generation of late-born neuronal subtypes across multiple classes of spinal cord neurons.

### Loss of miRNAs disrupts neural progenitor temporal patterning

Classical studies in insects and more recent observations in vertebrates suggest that chronological neuronal subtype specification is controlled by gene regulatory networks (GRNs) in neural progenitors that determine their temporal identity and define competence windows for the generation of specific neuronal progeny (*6*, *58–63*). We therefore tested whether loss of Dgcr8 affects temporal patterning of progenitors. DGE analysis showed a similar shift towards earlier temporal identities as observed in neurons. Genes associated with late progenitor states, including *Hopx*, *Nfib*, *Nfix*, and *Sall3*, astrocyte differentiation (*Atp1a2*, *Slc1a3*, *Slc6a11*, *Mt3*, *Aldoc*, *Ndrg2*) and oligodendroglial precursor cell (OPC) identity (*Olig1*, *Olig2*, *Sox10*, *Bcas1*) were downregulated. Conversely, mutant progenitors showed increased expression of genes characteristic for early progenitors, including *Prtg*, *Zbtb16*, *Nr6a1*, and *Trim71* (Fig. 4A). GO enrichment analysis revealed a clear separation of biological processes affected by loss of Dgrc8. Genes upregulated in mutant progenitors were enriched for pathways associated with cell cycle progression, DNA replication, and mitotic division (Fig. 4B), indicating a persistence of proliferative programs in the Dgcr8 cKO embryos, in agreement with the observed increase in mitotic activity in these embryos (Fig. S2 I, K). In contrast, downregulated genes were enriched for gliogenesis and neuronal differentiation-related functions, including synapse assembly and cognitive processes, suggesting a failure to properly engage maturation and functional specialization programs (Fig. 4C). Together, these findings suggest that loss of Dgcr8 impairs the normal temporal progression of neurons and progenitors, resulting in the persistence of early developmental programs and reduced acquisition of late-stage identities.

**Fig. 4.**
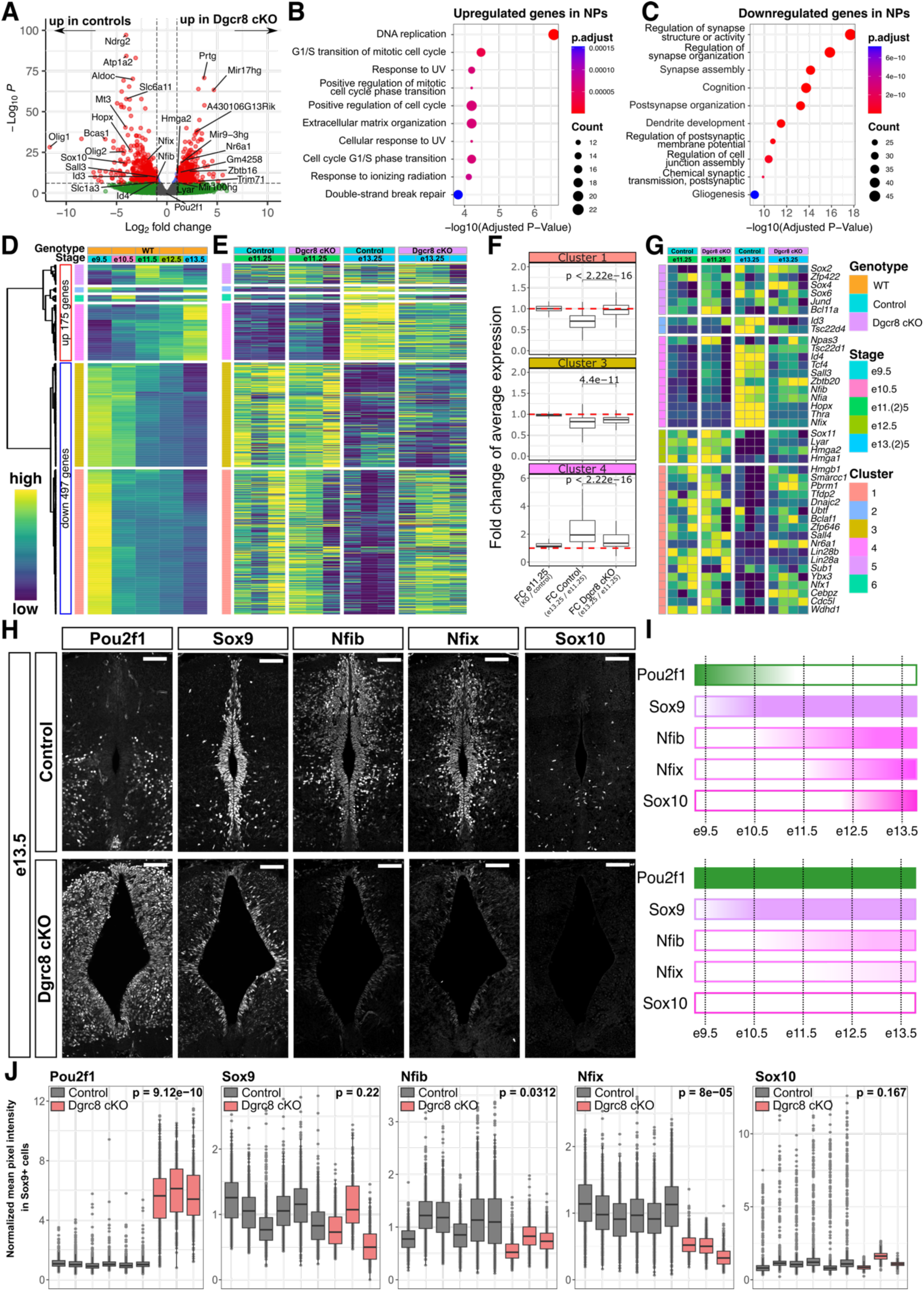
Temporal patterning of neural progenitors is perturbed in Dgcr8 cKO embryos. **(A)** DGE analysis of e13.25 neural progenitors reveals down-regulation of late and maintenance of early progenitor markers in Dgcr8 cKO embryos. **(B,C)** GO term associated with transcripts up-(B) and down-regulated (C) in progenitors in the e13.25 scRNAseq dataset **(D)** Differential gene expression analysis using scRNAseq from spinal cord neural progenitors (*8*, *10*) identifies 672 genes that are differentially expressed during the neurogenic period. Further segregation into 6 gene clusters with different dynamics is denoted by the dendrogram on the right. Heatmap shows log-scaled and z-scored gene expression values for each gene. **(E)** Analysis of the 672 differentially expressed genes in neural progenitors from the e11.25 and e13.25 scRNAseq datasets. **(F)** Average fold change for genes in clusters 1,3 and 4, when compared between Dgcr8 cKO and control embryos at e11.25 (left), control embryos at e13.25 and e11.25 (middle) and Dgcr8 cKO embryos at e13.25 and e11.25. **(G)** Expression dynamics of TFs included in the 672 genes in E. **(H)** e13.25 cervical spinal cord sections of control (top) and Dgcr8 cKO embryos stained for Pou2f1, Sox9, Nfib, Nfix, and Sox10. **(I)** Schematic indicating expression dynamics of the TFs in H in control (top) and Dgcr8 cKO embryos (bottom). **(J)** Quantification of normalized intensities for the indicated markers in Sox9+ neural progenitors from different control and Dgcr8 cKO embryos. Scale bars = 150 μm, Statistical significance in F was assessed by paired Wilcoxon signed rank test, statistical significance in J was assessed by applying a linear mixed effects model with genotype as fixed effect and embryo and litter id as random effects. p-values are integrated on the graphs.

To test this hypothesis, we sought to place the transcriptional changes observed in *Dgcr8* mutants within the broader context of gene expression dynamics during spinal cord development. To this end, we leveraged a scRNAseq dataset of the developing spinal cord (*10*). Identification of genes exhibiting broadly conserved temporal regulation during the neurogenic period in progenitors, as defined previously (*8*), yielded 672 genes, including 41 TFs, with 497 genes showing progressive downregulation and 175 genes being upregulated during development. This set was slightly larger than previously reported, likely reflecting updates in Seurat-based DGE analysis workflows; however, hierarchical clustering, heatmap structure, and identified TFs closely recapitulated the previously described patterns (*8*) (Fig. 4D-G). Using hierarchical clustering, we identified different expression dynamics of these genes. Genes in clusters 1 and 3 were sequentially downregulated after e9.5 or e10.5, respectively, genes in clusters 4 were induced during the neurogenic period, genes in cluster 5 peaked in the middle of the neurogenic period, and genes in clusters 2 and 6 were upregulated at the beginning and the end of the neurogenic period (Fig. 4D). We next analyzed the expression dynamics of these genes in the scRNAseq data from e11.25 and e13.25 control and Dgcr8 cKO embryos, focusing on downregulated (clusters 1 and 3) and induced genes (cluster 4). At e11.25, genes in these clusters were expressed at similar levels in control and Dgcr8 cKO embryos and, consistently, the fold change differences of the aggregated expression levels of the genes in these clusters were close to 1 (Fig. 4E-F). In contrast, at e13.25 genes in cluster 3 and in particular in cluster1 were much less repressed in the Dgcr8 cKO embryos, and genes in cluster 4 were significantly less induced. Many TF-encoding transcripts included in these clusters followed similar dynamics (Fig. 4G). Collectively, these results suggest miRNAs repress broad aspects of the early progenitor identity program. Notably, genes included in cluster 1, which are typically down-regulated between e9.5 and e10.5 were stronger affected by these changes than cluster 3 genes, which only disappear after e10.5 (Fig. 4D-F), suggesting that miRNAs contribute in particular to the down-regulation of the e9.5 progenitor transcriptional program.

To test this hypothesis, we examined the expression dynamics of progenitor temporal identity markers in control and Dgcr8 cKO embryos between e9.5 and e13.5. Because our previous analysis had identified expanded Pou2f1 expression in the ventricular zone of e13.5 Dgcr8 cKO sections (Fig. 2A, D), we speculated that Pou2f1 is expressed in early progenitors in the spinal cord and subsequently down-regulated as progenitors switch to later temporal identities, and that this down-regulation does not occur in the Dgcr8 cKO embryos. We therefore analyzed the expression of Pou2f1 in combination with the late temporal identity factors Sox9, Nfib and Nfix, which are sequentially induced in progenitors during the neurogenic period and play important roles in the transition from neurogenesis to gliogenesis, and the OPC marker Sox10 (*64–67*). These markers were indeed sequentially expressed in spinal cord progenitors in control embryos - Pou2f1 was expressed at e9.5, but by e11.5 its expression was largely extinguished in progenitors, while maintained in floor and roof plate cells and early-born neurons. Sox9 was induced at e9.5, Nfib at e11.5, and expression of Nfix in progenitors and Sox10 in OPCs emerged between e11.5 and e13.5 (Fig. S5A-B, Fig. 4H). In Dgcr8 cKO embryos, Pou2f1 levels were detectable in progenitors at e11.5, albeit reduced compared to the levels detected in neurons and floor plate cells (Fig. S5B). Notably, its expression in neurons was also expanded. In control embryos, Pou2f1 was mostly confined to neurons in the ventral half of the spinal cord, while it was present in neurons along the entire DV axis in Dgcr8 cKO embryos (Fig. S5B). These observations are consistent with the results of the EdU-birth-dating that Pou2f1+ neurons continue to be produced at e11.5 in the Dgcr8 cKO embryos (Fig. 2). At e13.5, Pou2f1 levels were increased in progenitors and reached similar levels as in neurons and floor plate cells. Maintained expression of Pou2f1 was accompanied by reduced expression of the late markers Nfib and Nfix, while expression of Sox10 was essentially undetectable (Fig. 4H-J). In contrast, Sox9 levels were not significantly altered, suggesting most progenitors in Dgcr8 cKO embryos acquire a Sox9, Pou2f1 double-positive identity characteristic for early neurogenic progenitors in the spinal cord (Fig. 4H-J).

Collectively, these data further corroborate the finding that temporal patterning of progenitors and the transition to late identity stages is severely disrupted in Dgcr8 cKO embryos. These data moreover reveal Pou2f1 as a novel tTF in the spinal cord that is expressed in early progenitors and subsequently maintained in early-born neurons. This finding is consistent with a model in which Pou2f1 regulates temporal patterning of progenitors and neurons in the spinal cord, similar to its role in controlling the temporal window for cone generation in the retina (*22*).

### Evolutionary conserved targeting of the early temporal program in progenitors and neurons by let-7 and miR-9

miRNAs act primarily through transcript destabilization and translational repression by sequence specific binding to the 3’-UTRs of target mRNAs, typically mediated by the miRNA seed region at nucleotides 2-7 or 2-8 from the 5′ end of the miRNA (*24*, *25*, *27*). To identify candidate miRNAs underlying the observed temporal patterning defects, we performed miRNA seed enrichment analysis using enrichMir (*68*) on transcripts predicted to be upregulated in progenitors or neurons (p < 1 × 10⁻⁶; log2-fold change > 1) in the Dgcr8 cKO mutants at e13.25. We reasoned that these are the most likely direct targets released from miRNA-mediated repression. let-7-5p and miR-9-5p seeds were statistically enriched in the 3’-UTRs of upregulated transcripts in progenitors and neurons, while miR-124-3p.1 and miR-124-3p.2 seeds were enriched only in neurons (Fig. 5A). These observations are consistent with the neuron-specific expression and function of the miR-124 family (*69*, *70*), and reveal let-7 and miR-9 as candidates for regulating temporal patterning of progenitors and neurons in the spinal cord.

**Fig. 5.**
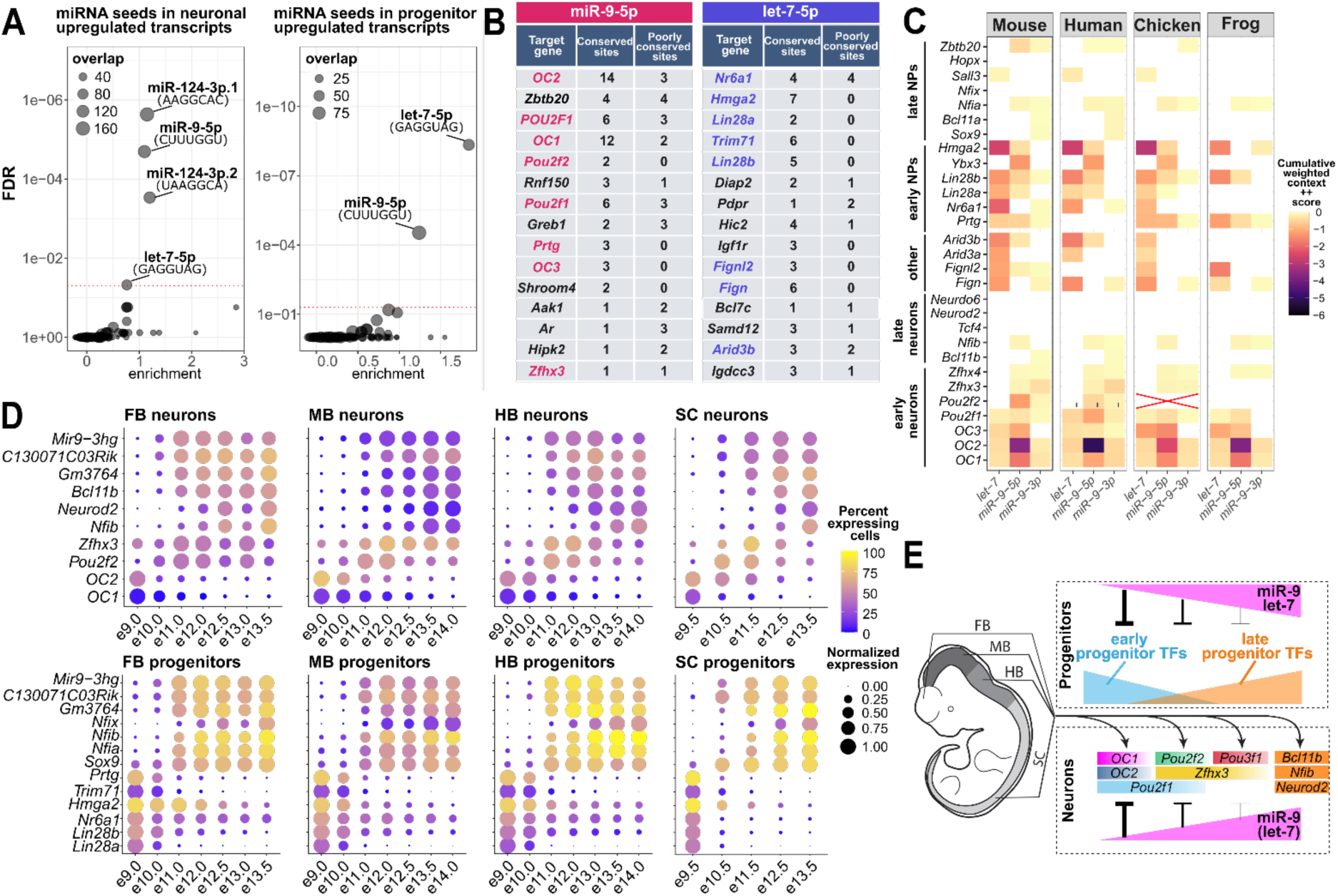
Computational analysis identifies miR-9 and let-7 as candidate miRNAs involved in temporal patterning. **(A)** miRNA seed enrichment analyses on transcripts upregulated in neurons (left) and progenitors (right) from Dgcr8 cKO embryos. miR-9-5p and let-7-5p seeds are enriched in progenitors and neurons, miR-124-3p.1 and miR-124-3p.2 seeds are enriched only in neurons. **(B)** Top-15 miR-9-5p (left) and let-7-5p (right) targets in mouse predicted by Targetscan. Colored font highlights transcripts upregulated in progenitors and neurons from Dgcr8 cKO embryos. **(C)** Heatmap indicating the number of predicted miRNA target sites in the 3’-UTRs of mRNAs encoding early and late progenitor and neuronal markers predicted by Targetscan. Heatmap on the left indicated the observed log2 fold-change observed in the scRNAseq data from the Dgcr8 cKO embryos **(D)** Expression dynamics of miR-9 host genes in scRNAseq data from the forebrain, midbrain, hindbrain, and spinal cord (from left to right) in relation to other early and late neuron (top) and progenitor (bottom) markers. **(E)** Rising miR-9 and let-7 levels in progenitors progressively repress early progenitor markers and thereby promote the switch from early to late progenitor identities. miR-9 acts additionally in neurons by repressing early neuronal tTFs such as OC TFs, which have a large number of miR-9 target sites in their 3’-UTRs. Later tTFs have fewer miR-9 target sites in their 3-UTRs and are thus progressively less repressed by miR-9.

Post-transcriptional regulation of the early progenitor markers *Lin28a*, *Lin28b* and *Trim71* by let-7 is evolutionary highly conserved (*71–74*). Similarly, *Lin28a* and *OC* transcripts have been previously identified as miR-9 targets in different developmental and disease contexts (*41*, *45*, *75–77*). We therefore speculated that miR-9 and let-7 directly regulate other early progenitor and neuronal identity markers. TargetScan analysis identified numerous early neuronal tTFs, including all three *OC* TFs, *Pou2f1, Pou2f2*, *Zfhx3* and *Zfhx4* as miR-9-5p targets (Fig. 5B) (*78*, *79*). Consistent with recent observations (*76*), we confirmed in dual-luciferase assays that miR-9 represses reporter constructs fused to *OC2* and *Pou2f1* 3’-UTR fragments, providing experimental evidence for the functionality of these sites (Fig. S6A-B). Constructs fused to different fragments of the *Pou2f2* and *Zfhx3* 3’-UTRs, which each contained only one miR-9-5p binding site (Fig. S6A), were only repressed moderately or not at all (Fig. S6B), consistent with previous observations that cooperativity among multiple target sites is required for efficient mRNA silencing (*80*). Notably, the number of miR-9-5p seed sequences predicted by Targetscan and the expected repression, indicated by Targetscan’s Cumulative weighted context ++ score (CWCS) (*78*), correlated with the chronological order, in which neurons expressing these tTFs are generated. The 3′UTRs of *OC1* and *OC2*, which define the earliest neuronal identities, contained 12 and 14 evolutionarily conserved and 2 and 3 poorly conserved miR-9-5p binding sites, respectively, while *Pou2f1*, *Pou2f2*, *Zfhx3* and *Zfhx4* harbored 6, 2, 1 and 1 evolutionary conserved sites (Fig. 5B-C, Fig. S6C-D). *Pou2f1* and *Zfhx3* contained moreover 3 and 1 poorly conserved miR-9-5p sites (Fig. 5B-C, Fig. S6C-D). Consistently, the CWCSs of early tTFs, in particular *OC1* and *OC2*, were more negative than for later tTFs, predicting stronger repression of early target genes by miR-9. Other miR-9-5p targets included the early progenitor markers *Lin28a*, *Lin28b*, *Trim71*, *Prtg*, and *Ybx3*, as well as *Fignl2*, which was strongly upregulated in neurons from Dgcr8 cKO animals (Fig. 5B-C, Fig. S6C-D). Moreover, many of these transcripts contained seed sequences for miR-9-3p (Fig. 5C, Fig. S6D). Similarly, multiple early progenitor markers including *Nr6a1*, *Hmga2*, *Lin28a*, *Lin28b, Prtg*, *Pou2f1 and Trim71,* as well as *Fign*, *Fignl2, Arid3a* and *Arid3b*, which were up-regulated in Dgcr8 cKO neurons, were predicted let-7-5p targets (Fig. 5B-C, Fig. S6C-D). Moreover, miR-9-5p and let-7-5p target many of these transcripts in other vertebrates including human, chicken, frog, and fish, suggesting deep evolutionary conservation of the let-7 and miR-9-mediated antagonism of early temporal identity programs of progenitors and neurons in vertebrates (Fig. 5B-C, Fig. S6C-D). The striking correlation between the chronological birth order of neurons expressing different tTFs and the number of miR-9 seed sequences, and therefore the expected repression of these transcripts by miR-9, raises moreover the exciting possibility that miR-9 levels define the competence windows for neuronal tTF expression.

### Characterization of let-7 and miR-9 expression

Previous work and our scRNAseq analysis demonstrated that let-7 and miR-9 are present in the embryonic spinal cord (*44*, *45*, *81*); however, whether the expression dynamics of these miRNAs correlate with the sequential transition from early to late progenitors and how their expression patterns relate to the expression patterns of neuronal and progenitor tTFs is currently unclear. Because mature miRNAs are typically not detected by conventional scRNAseq and RNAseq, we analyzed the expression of pri-miRNA transcripts in relation to early and late temporal identity markers of progenitors and neurons in embryonic scRNAseq time-course datasets spanning the spinal cord, hindbrain, midbrain and forebrain (*10*, *82*). Across all CNS regions, expression of the miR-9 family pri-miRNAs *C130071C03Rik*, *Mir9-3hg* and *Gm3764* progressively increased during development in progenitors and neurons (Fig. 5D). This increase coincided with the downregulation of early temporal markers and the emergence of intermediate and late temporal identities. Correlation-based analysis moreover revealed that *Gm3764*, *C130071C03Rik* and *Mir9-3hg* were among the transcripts most strongly correlated with the expression of the late temporal marker *Nfib* in progenitors and neurons from all analyzed regions (Fig. S6E, Fig. S7A), and most anti-correlated with the early tTF *OC2* in neurons (Fig. S6F). Unlike for the miR-9 pri-miRNAs, we did not detect expression of let-7 pri-miRNAs such as *Mir99ahg* or *Mir100hg* in any dataset. Together, these data suggest that miR-9 is progressively induced in progenitors throughout the developing nervous system and that its levels in neurons correlate with temporal identity.

We next sought to validate these dynamics experimentally. To this end, we first analyzed let-7 and miR-9 expression dynamics in in-vitro differentiations of dorsal and ventral spinal cord-like neurons from mouse embryonic stem cells (ESCs) that recapitulate temporal patterning of progenitors and neurons (Fig. S7B-D) (*8*). RT-qPCR analysis revealed progressive upregulation of miR-9-5p, miR-9-3p, let-7a and let-7c in dorsal and ventral differentiation paradigms, consistent with the expression dynamics observed in-vivo. We next tested whether the progressive increase in miR-9 is mediated by the sequential activation of miR-9 host genes. Analysis of the expression dynamics of the three miR-9 pri-miRNAs *Gm3764*, *C130071C03Rik* and *Mir9-3hg* revealed distinct temporal expression profiles. *C130071C03Rik* and *Mir9-3hg* sharply increased between differentiation day (D) 5 and D7, whereas induction of *Gm3764* was comparatively delayed, occurring between D7 and D9 (Fig. S7E). These temporally distinct activation dynamics coincide with the down-regulation of the early temporal program and the emergence of late progenitor and neuronal identities in this system (Fig. S7B) (*8*, *60*), further corroborating the hypothesis that the progressive accumulation of miR-9, driven by the sequential activation of miR-9 host genes (*83*), contributes to the temporal maturation of progenitors and neurons.

We next aimed to validate these observations in the embryonic spinal cord. To this end, we detected let-7 and miR-9 in combination with *Pou2f2* and *Nfib* mRNAs in e13.5 spinal cord sections from control and Dgcr8 cKO embryos by RNAscope. In control embryos, let-7 and miR-9 levels were anti-correlated with *Pou2f2* expression and elevated in medial *Nfib+* progenitors and neurons (Fig. 6A-D). Signals for both miRNAs were markedly reduced in spinal cords of *Dgcr8* cKO embryos, confirming the specificity of both signals for the mature forms of both miRNAs (Fig. 6A-D). The reduction in miRNA levels was accompanied by loss of *Nfib* expression and a corresponding expansion of *Pou2f2* (Fig. 6A-D). Strikingly, both miRNA patterns closely mimicked the previously established pattern of neuronal birth-dates in the spinal cord, with low levels in the lateral parts of the ventral horns, where most early-born spinal cord neurons reside, and higher levels in the medial parts of the ventral spinal cord and the dorsal horns. miR-9 RNAscope in combination with immunofluorescence for different tTFs further confirmed anti-correlated expression of early tTFs, such as OC2, Pou2f2 and Zfhx3, and miR-9, while late tTFs such as Neurod2 and Nfib were confined to regions with high miR-9 levels (Fig. S7F).

**Fig. 6.**
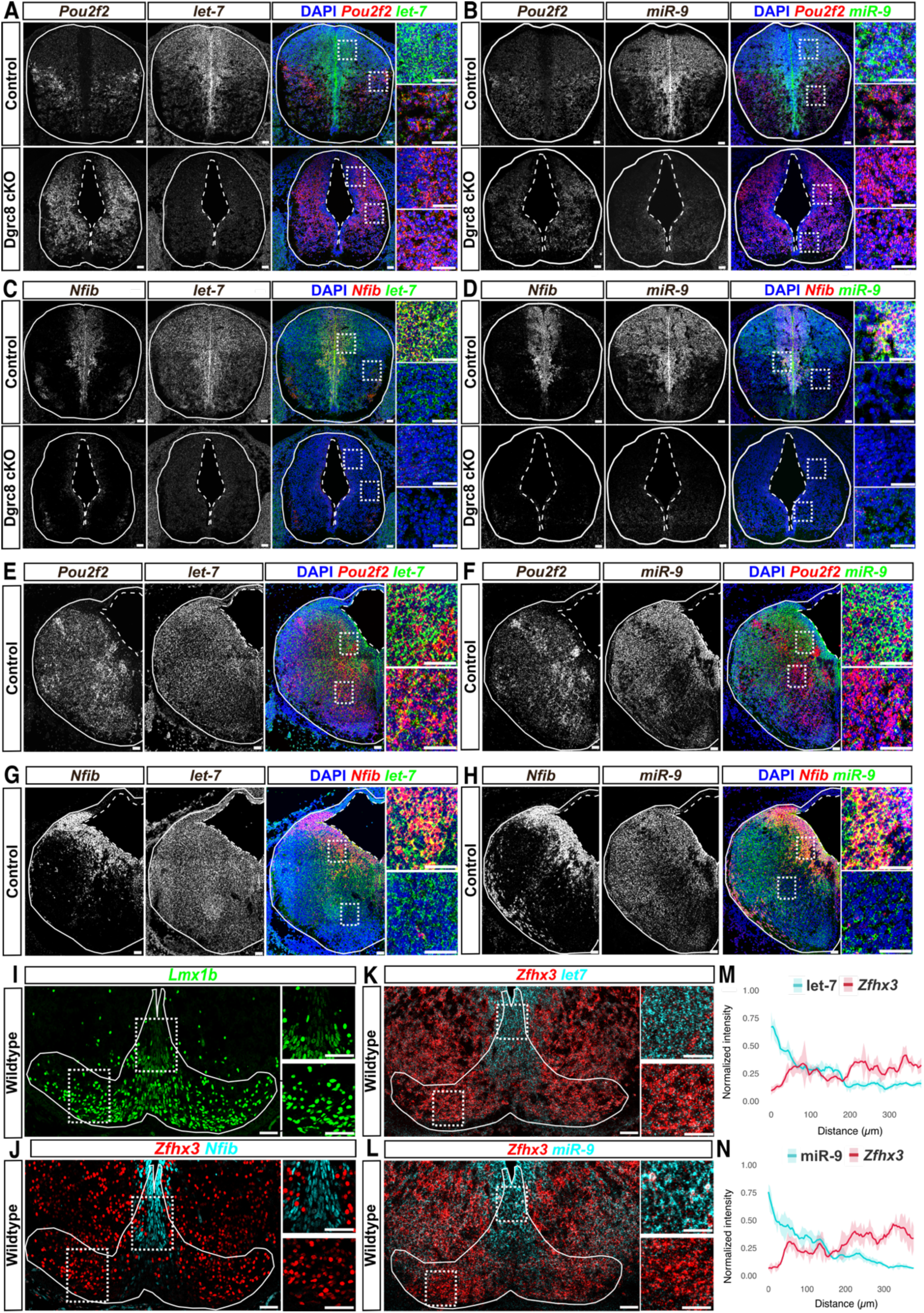
miR-9 and let-7 expression correlates with the chronology of neuronal subtype specification. **(A-D)** RNAscope of e13.5 cervical spinal cord sections from control (top) and Dgcr8 cKO (bottom) embryos for let-7 (A, C) or miR-9 (B, D) in combination with *Pou2f2* (A, B) or *Nfib* (C, D). let-7 and miR-9 expression is low in *Pou2f2*+ areas and closely follows the described patterns of neuronal birth in the spinal cord. Levels of both miRNAs and *Nfib* are reduced and *Pou2f2* levels are elevated in sections from Dgcr8 cKO embryos. **(E-H)** RNAscope of e13.5 hindbrain sections from control embryos for let-7 (E, G) or miR-9 (F, H) in combination with *Pou2f2* (E, F) or *Nfib* (G, H). As in the spinal cord, let-7 and miR-9 expression is low in *Pou2f2*+ areas. **(I, J)** mDA neurons, identified by Lmx1b expression (I), can be partitioned into an early-born lateral Zfhx3+ and late-born medial Nfib+ neurons (J). **(K, L)** RNAscope of e13.5 ventral midbrain sections reveals anti-correlated expression of let-7 and *Zfhx3* (K) and miR-9 and *Zfhx3* (L). Levels of both miRNAs are high in medial areas, where Nfib+ progenitors and neurons reside, and low in lateral Zfhx3+ areas. **(M, N)** let-7 and *Zfhx3* (M) or miR-9 and *Zfhx3* (N) RNAscope levels, quantified from medial to lateral along the dorsal edge of the Lmx1b expression domain. Scale bars = 50 µm

To test if similar (anti-)correlations are present in other parts of the CNS, we analyzed let-7 and miR-9 expression in combination with *Pou2f2* and *Nfib* in e13.5 hindbrain sections (Fig. 6E-H), and in combination with *Zfhx3* in midbrain dopaminergic (mDA) neurons, which are also partitioned into an early Zfhx3+ and a late Nfib+ subset (*8*, *84*) (Fig. 6I-N). Consistent with our observations in the spinal cord, miR-9 and let-7 were low in *Pou2f2+* neurons in the hindbrain and *Zfhx3+* neurons in mDA neurons and elevated in late-born Nfib+ progenitors and neurons in both areas (Fig. 6E-N). Together, these observations provide experimental evidence for the presence of temporal miRNA gradients that restrict the expression of early tTFs in large swathes of the developing CNS.

### Minor effects of let-7 perturbations on temporal patterning

We next aimed to test if manipulating miR-9 or let-7 levels and expression dynamics perturbs temporal patterning in an in-vivo context. To this end, we focused on the developing chicken spinal cord, a well-established model amenable to transient gain and loss-of function approaches by electroporation (*85*, *86*). Analysis of tTF expression by immunofluorescence revealed moreover largely conserved temporal patterning of the mouse and chicken embryonic spinal cord (Fig. S8), consistent with recent observations by scRNAseq (*87*).

To analyze whether perturbing let-7 levels affects temporal patterning in the developing spinal cord, we electroporated expression constructs for let-7c and nuclear DsRed into chicken embryos at HH15-16 and assessed the consequences by staining for the early tTFs Pou2f1 and Zfhx3, and the late marker Neurod2 at HH25. We chose let-7c because it is derived from the *Mir99ahg* pri-miRNA that was up-regulated in the spinal cord of Dgcr8 cKO embryos, suggesting it is endogenously expressed in the embryonic spinal cord, and its mature sequence is completely preserved between mouse and chicken. Ectopic expression of let-7c did not alter temporal patterning, as judged by the expression of Pou2f1, Zfhx3, and Neurod2 (Fig. S9A-B). We next asked whether overexpression of Lin28a, which inhibits maturation and thereby antagonizes activity of let-7 miRNAs (*88–90*), affects temporal identity. Similar to let-7, ectopic Lin28a expression had little impact on overall temporal patterning, although we consistently observed a small population of faint ectopic dorsal Pou2f1+ cells within the electroporated domain. This induction was markedly weaker than the endogenous Pou2f1 expression observed in neurons at this stage (Fig. S9C-D). Collectively, these results suggest a minor role for let-7 in controlling temporal patterning of progenitors and neurons in the embryonic spinal cord.

### Perturbing miR-9 expression dynamics affects chronological neuronal subtype generation

In contrast, ectopic expression of miR-9 resulted in an approximately 25% reduction in the number of neurons expressing the early tTFs Pou2f1, Zfhx3, and Zfhx4 (Fig. 7A-B; Fig. S9E-F). Similarly, miR-9 expression significantly reduced the number of neurons expressing the early neuronal tTF Meis2 (*11*) by approximately 10% (Fig. S9E-F). Consistent with a shift in temporal identity, miR-9 expression increased the number of Bcl11b+ neurons by approximately 20% (Fig. 7A-B). Even more strikingly, Neurod2, which is normally undetectable at this stage, was robustly induced following miR-9 expression (Fig. 7A,C). Importantly, the effects of miR-9 expression appeared largely cell-autonomous. Very few DsRed+ cells expressed the early tTFs Pou2f1 or Zfhx3 (Fig. 7A; Fig. S9E-F), and if they did, levels of these TFs were reduced in DsRed+ neurons (Fig. 7D). In contrast, the majority of Bcl11b+ and all Neurod2+ neurons expressed DsRed (Fig. 7A), with significantly higher Bcl11b and Neurod2 signal intensities detected in DsRed+ neurons (Fig. 7D). These effects were specific to miR-9, as electroporation of the same construct lacking the miR-9 sequence did not alter tTF expression (Fig. S9G).

**Fig. 7.**
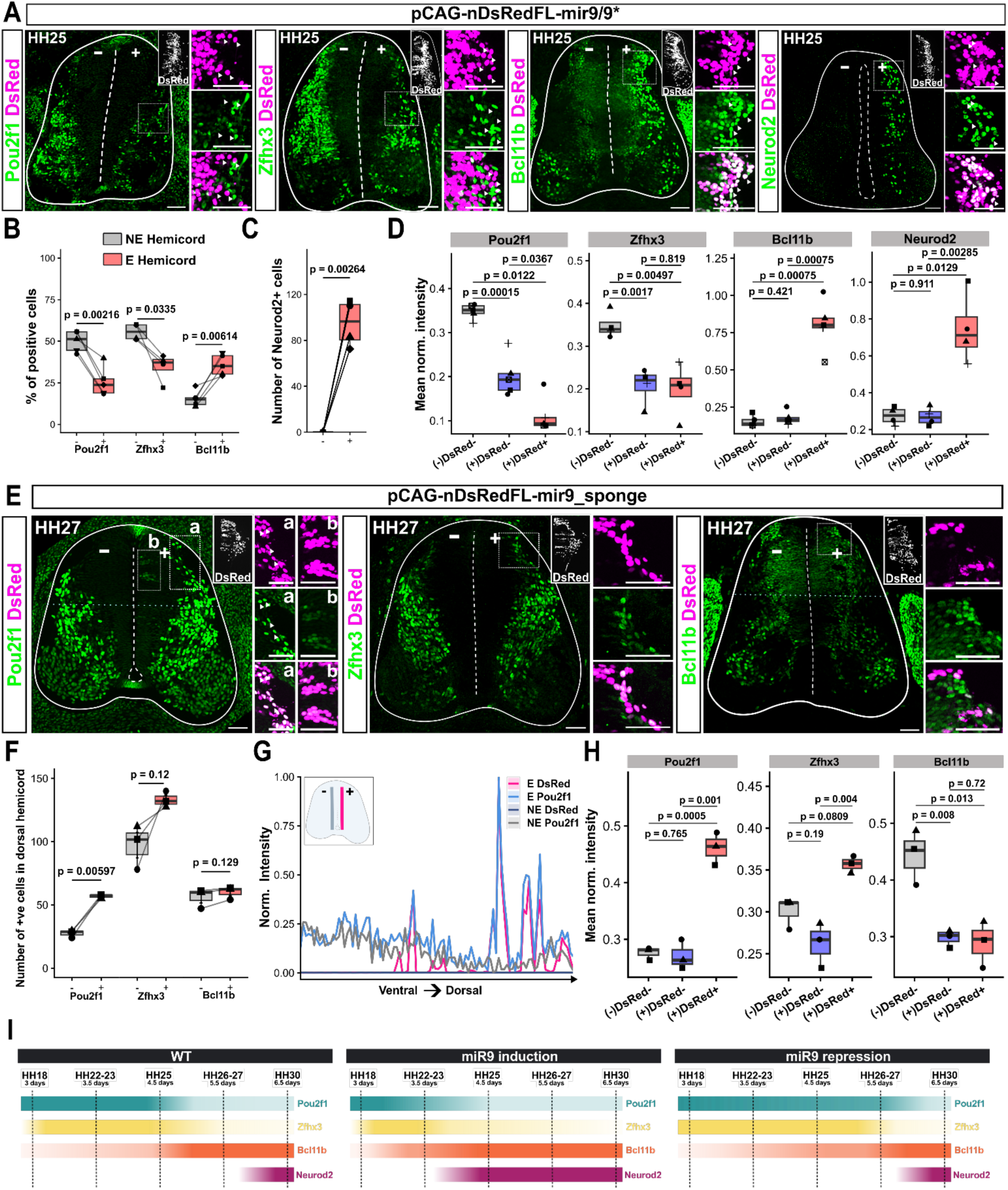
miR-9 expression perturbations alter temporal patterning of the embryonic chicken spinal cord. **(A)** Ectopic miR-9 expression, indicated by presence of nuclear DsRed (nDsRed), represses the early tTFs Pou2f1 and Zfhx3 and induces the late tTFs Bcl11b and Neurod2 in the chicken spinal cord at HH25. **(B, C)** Percentage (B) or number (C) of neurons expressing the indicated markers on the electroporated (E, +) and non-electroporated (NE, -) site of the spinal cord. **(D)** Average normalized intensity of the indicated markers on the non-electroporated site (-), and in non-electroporated [DsRed-] and electroporated cells [DsRed+] on the electroporated site (+). Late markers Bcl11b and Neurod2 are induced cell-autonomously in (+)DsRed+ cells. **(E)** Expression of a nDsRed-miR-9 sponge construct promotes expression of early tTFs Pou2f1 and Zfhx3 and represses Bcl11b in the chicken spinal cord at HH27. **(F)** Number of neurons in the dorsal half of the spinal cord on the non-electroporated (-) and electroporated (+) site expressing the indicated markers. **(G)** Intensity histograms of the electroporated (E) and non-electroporated (NE) sites for DsRed and Pou2f1 of the spinal cord shown in E. Expression of the miR-9 sponge, indicated by nDsRed expression, maintains Pou2f1 expression in progenitors. **(H)** Average normalized intensity of the indicated markers on the non-electroporated site (-), and in non-electroporated [DsRed-] and electroporated cells [DsRed+] on the electroporated site (+). Levels of early markers Pou2f1 and Zfhx3 in nDsRed+ cells are elevated. **(I)** Schematics indicating the phenotypic consequences of miR-9 expression perturbations. Compared to the wildtype (left), miR-9 induction causes precocious termination of Pou2f1 and Zfhx3 expression and induction of Neurod2 and Bcl11b (middle). In contrast, antagonizing miR-9 activity with a miR-9 sponge extends the window during which Pou2f1+ and Zfhx3+ neurons are generated and antagonizes Bcl11b expression in neurons. Scale bars = 50 µm. Statistical significance in B, C, and F (Pou2f1 and Bcl11b) was assessed using a paired t-test; in F (Zfhx3), using a Wilcoxon rank-sum test; and in D and H, using a Student’s t-test, Welch’s t-test, or Wilcoxon rank-sum test, as appropriate based on the normality and homoscedasticity of the data. p-values are indicated on the graphs.

To determine whether endogenous miR-9 is required for temporal progression, we next inhibited miR-9 activity using a miR-9 sponge containing 16 tandem binding sites inserted in the 3’-UTR of the DsRed expression vector. Inhibition of miR-9 resulted in a significant increase in both the number of Pou2f1+ cells and the intensity of Pou2f1 expression. This effect was particularly obvious in the dorsal spinal cord, where Pou2f1 is normally absent at this stage because of the ventral migration of neurons from most dorsal classes (Fig. 7E, F, H). Notably, ectopic Pou2f1 expression was also detected in progenitors and strongly correlated with DsRed expression (Fig. 7E-G). We also observed a trend toward increased Zfhx3 expression. Although the number of Zfhx3+ cells was not significantly altered, Zfhx3 signal intensity was significantly increased following miR-9 inhibition (Fig. 7E, F, H). Finally, DsRed and Bcl11b expression remained largely mutually exclusive, suggesting reduced Bcl11b induction upon miR-9 inhibition (Fig. 7E-F).

These results support a model in which miR-9 promotes temporal progression by enabling the transition from early to late temporal identities. Ectopic miR-9 expression prematurely represses early tTFs and promotes the expression of late markers, whereas miR-9 inhibition prolongs the expression windows of early tTFs (Fig. 7I). Collectively, these data establish miR-9 as major regulator of temporal identity in the spinal cord, whose expression dynamics dictate the sequential transition from early to late progenitor states and the chronologically coordinated generation of distinct temporal cohorts of neurons.

## Discussion

Our study identifies miRNAs as essential regulators of temporal patterning throughout the developing CNS by demonstrating that disruption of miRNA biogenesis through Dgcr8 deletion prevents the normal progression from early to late developmental identities. Across the spinal cord, hindbrain and midbrain, loss of Dgcr8 resulted in the persistence and prolonged production of early-born neuronal populations at the expense of late-born neurons, revealing a conserved requirement for miRNAs in advancing the temporal program. These defects could not be explained solely by altered proliferation, apoptosis or tissue growth, but instead reflected an extension of the developmental windows during which early temporal identities are generated. Single-cell transcriptomic analyses further showed that, although broad dorsoventral neuronal identities were largely preserved, both neurons and progenitors exhibited a widespread retention of early transcriptional programs, impaired activation of late differentiation and maturation genes, and delayed developmental progression. Integrative computational analyses identified miR-9 and let-7 family members as the strongest candidate regulators of these processes, with expression dynamics that closely mirror temporal progression across the embryonic CNS. Functional and molecular experiments further established miR-9 as a key component of the temporal patterning network that directly represses early tTFs, promotes the timely acquisition of late neuronal identities, and drives temporal progression in-vivo. Together, our findings support a model in which the progressive accumulation of miRNAs, particularly miR-9, acts as a conserved molecular timer that coordinates the transition from early to late developmental states in neural progenitors and neurons, thereby ensuring the orderly generation of neuronal diversity during vertebrate CNS development.

### Comparison with temporal patterning in other developmental contexts

A central question raised by our findings is how miRNAs interface with previously described temporal patterning mechanisms in other regions of the developing nervous system. In this regard, our results parallel observations in the developing retina, where miRNAs have also emerged as critical regulators of temporal identity transitions (*41*, *50*, *91*). More specifically, miR-9-2 regulates timing of retinal progenitor cell competence and maintenance of Müller glial identity (*42*), while let-7 has been shown to regulate cell cycle dynamics (*92*). Our findings indicate that this regulation is not restricted to retinal development, but represents a broader principle governing temporal patterning across the CNS (Fig. 1, Fig. S1B-F), consistent with the previous identification of temporal programs of neurons and progenitors that apply throughout the developing CNS (*8*). The conservation of miRNA candidates and their expression dynamics (Fig. 6, Fig. S7) suggests that miRNAs constitute a general molecular mechanism by which neural tissues progressively restrict early developmental programs and acquire later identities. However, our results also reveal interesting differences to other brain regions. In the cortex, miR-9 and let-7 form temporally opposite and functionally antagonistic gradients, with ectopic miR-9 selectively specifying ‘early’ cortical layer V neurons, while let-7 promotes acquisition of superficial layer identities typically generated later during development (*39*). These differences may be due to cortical neurons transcriptionally resembling late-born neurons in other parts of the CNS and their overwhelming generation from progenitors that share many features with late neuronal or even glial progenitors in other parts of the nervous system (*8*). Further comparison of the transcripts regulated by the respective miRNAs in the different areas is required to understand how the combinatorial activities and expression dynamics of both miRNAs temporally pattern different parts of the embryonic CNS.

In *Drosophila*, temporal patterning is controlled by the sequential expression of temporal identity factors in progenitors, that instruct the generation of specific neuronal progeny at defined temporal windows during development (*58*, *59*, *63*). In the ventral nerve cord, a temporal TF expression cascade of Hunchback ◊ Krüppel ◊ Nubbin/Pdm2 ◊ Castor ◊ Grainyhead directs the generation of distinct neuronal subtypes in essentially all neuroblast lineages. These TFs are typically expressed first in neuroblasts and their expression is subsequently maintained in the specific temporal cohorts of neurons generated from the neuroblasts during their expression windows (*6*, *58*, *93*). Vertebrate orthologues of these core tTFs have been implicated in temporal patterning of the cortex and retina (*22*, *94–96*). Our current work identifies Pou2f1 as tTF in the spinal cord that, similar to its *Drosophila* orthologues Nubbin/Pdm2, is first expressed in progenitors and subsequently maintained in early born neurons (Fig. 4H-J, Fig. S5). Similar expression dynamics likely apply to the developing mid- and hindbrain, and have been documented in the retina (*22*). Pou2f1 may thus constitute a general and evolutionary deeply-conserved tTF involved in the specification of early neuronal identities in large parts of the CNS and animal kingdom. Whether orthologues of the other core temporal TFs initially identified in *Drosophila* are also involved in chronological neuronal subtype generation in the spinal cord is currently unclear and will be an exciting prospect for future investigation. Interestingly, among the transcripts most-upregulated in the e13.25 Dgcr8 cKO neurons were the Krüppel-like factors *Klf3* and *Klf7* (Fig. S4A), suggesting a potential role for both TFs in the establishment of early neuronal identities in the spinal cord.

Our findings moreover suggest that the molecular logic, which underlies temporal patterning differs between vertebrates and insects. In *Drosophila*, progression through the temporal sequence is largely mediated intrinsically by the tTFs cross-regulating each other’s expression (*58*, *59*). Once activated *Drosophila* neuroblasts will progress through large parts of the sequence independently of external factors, such as cell division and overall embryo development. In contrast, our results suggest that early TFs are dominant over later TFs and that miRNA-mediated repression of early identity factors is required to advance the temporal program of progenitors and neurons (Fig. 2, Fig. 4). These observations are consistent with the finding that within the OC2 → Pou2f2 → Pou3f1 sequence that chronologically partitions early motor and excitatory neurons in the spinal cord, early tTFs repress the subsequent identities in some instances, but later tTFs do not restrict the temporal competence windows during which earlier neuronal subtypes are specified (*11*). Our data suggest that this effect is at least partially mediated by rising miRNA levels, in particular of miR-9, during development that first inhibit OC TFs and at higher levels Pou2f1 and Pou2f2. This model is further supported by our observations that let-7 and miR-9 levels closely resemble the patterns of neuronal birth dates in the spinal cord (Fig. 6A-D), and that ectopic miR-9 expression represses Pou2f1 and prematurely induces the late temporal markers Bcl11b and Neurod2 (Fig. 7, Fig. S7B-E). These observations are moreover consistent with previous observations in motor neurons that linked progressively increasing miR-9 levels to the sequential generation of distinct motor neuron subtypes (*44*, *45*).

### microRNAs control temporal patterning by targeting progenitor and neuronal transcripts

Another question that is currently unresolved is whether the miRNA-mediated regulation of temporal identity predominantly occurs in progenitors, neurons, or both? Studies in *Drosophila* highlighted the importance of temporal patterning specification in stem cells and neuroblasts, with the temporal identity of the neuroblasts largely determining the temporal identity of their neuronal progeny (*6*, *58*, *63*, *93*). Similar observations have been made in the vertebrate cortex and retina, where the specification of distinct laminar identities by temporal miRNAs has been mostly ascribed to their activity in progenitors (*39*, *41*, *50*). In contrast, recent characterization of progenitor temporal patterning of multiple regions of the vertebrate CNS suggested the existence of only two clearly defined transcriptional states that each appear to give rise to multiple sequentially generated subtypes of neurons (*97*). For example, spinal neurons generated from early progenitors initially co-express OC2, Pou2f2, and Pou3f1 at low levels, before eventually resolving to a single-positive state in a chronologically coordinated manner (*11*). Early progenitors thus appear to be competent to generate the neuronal identities associated with all these TFs and the precise choice is only made in a stage-dependent manner at the post-mitotic level. Similarly, late-stage progenitors, characterized by the expression of Sox9 and NFI TFs in the spinal cord, sequentially generate multiple distinct cohorts of excitatory and inhibitory neurons that populate the distinct laminae of the dorsal horns (*13*). While our data do not rule out that additional temporal progenitor states exist, which give rise to specific temporal cohorts of neurons and that are potentially defined by the combinatorial expression of TFs, such as Nr6a1, Pou2f1, Sox9 and increasing NFI TF levels (*60*, *98*, *99*), our data are consistent with miRNAs acting in both progenitors and neurons to define temporal identity. First, miRNAs are required for the transition from the early to late progenitor state, as indicated by the persistent upregulation of early progenitor markers and failure to induce multiple aspects of a late progenitor identity, such as Nfib, Nfix and the gliogenic program, in Dgcr8 cKO embryos (Fig. 4, Fig. S5). Notably, multiple early progenitor markers such as Nr6a1, Prtg, Pou2f1 and Lin28b are targets of both miR-9 and let-7, suggesting that both miRNAs act together to promote this switch (Fig 5B-C). In particular miR-9 additionally targets multiple transcripts encoding neuronal tTFs, such as OC1, OC2, Pou2f1, Pou2f2 and Zfhx3, but for example also Foxp1 (*44*), which is only expressed in motor neurons in the early spinal cord (*100*). miR-9 levels in neurogenic cells and/or neurons may thus further refine, which specific neuronal temporal identities are generated from early and late progenitors by targeting global and neuronal subtype specific tTFs.

### Graded miRNA levels restrict the temporal identity of progenitors and neurons

The regulation of temporal patterning by rising miRNA levels conceptually resembles the affinity threshold model of morphogen signalling, except that the miRNAs do not form a gradient in space but in time and act at the post-transcriptional instead of the transcriptional level. Initially, low levels of miRNAs are only sufficient to repress transcripts with the highest affinity miRNA binding sites in their 3’-UTRs, similar to how low levels of morphogen signalling only activate target genes with more and high affinity binding sites for morphogen effectors far away from the source. As their levels rise, miRNAs are able to bind and thereby silence transcripts with fewer and progressively more suboptimal miRNA seed sequences in their 3’-UTRs, similar to how higher levels of morphogen effectors are able to bind and induce progressively less-optimal cis regulatory elements closer to the source. Of note, miRNA gradients create sharp gene expression domains and robustly position gene expression boundaries in a threshold-dependent manner during plant development (*101*, *102*), suggesting such a mechanism is generally feasible. Moreover, cross-regulatory interactions between the down-stream components of the miRNA-controlled GRN, such as those proposed between OC TFs, Pou2f2 and Pou3f1 (*11*), may similar to cross-repressive interactions between spatial TFs controlled by morphogens (*103*), further canalize the temporal GRN into discrete identities and confer robustness and hysteresis to the system. Components of the temporal program in progenitors and neurons may also actively regulate the effective levels of let-7 and miR-9, either by regulating the expression of the respective pri-miRNAs or interfering with let-7 and miR-9 biogenesis, for example by inducing or repressing Lin28a/b expression. Moreover, *OC2* mRNA, which contains an exceptionally long 3’-UTR with a high number of miR-9 seed sequences (*77*), has been proposed to act as a competing endogenous RNA to miR-9, that protects the transcriptional program induced by OC2 protein from miR-9 mediated repression. Such mechanisms may introduce further non-linearities into how the chronologically graded expression of pri-miRNAs is translated into effective mature miRNA concentrations, similar to how spatial target genes have been proposed to influence morphogen responsiveness to convert graded morphogen input into step-wise changes in gene expression (*104*). Thus, similar to spatial morphogen signalling, where the architecture of the downstream GRN determines the spatial and temporal response to graded morphogen input (*1*, *103*), temporally-graded miRNAs may serve as key regulators for the temporal patterning GRN, whose dynamics, determined by GRN architecture, convert levels and duration of miRNA input into discrete temporal phases of TF expression.

## Materials and Methods

### Animal welfare

Animal experiments were performed in accordance with the German and European animal welfare laws.

### Mouse lines

*Dgcr8^flox/flo^*^x^ (*49*), *Brn4-Cre* (also known as Tg(Pou3f4)-Cre) (*48*) and *Rosa26-EYFP* (*105*) mouse lines were used to perform the experiments in this work. The *Dgcr8^flox/flo^*^x^ line was crossed with the *Brn4-Cre* transgenic mice bearing heterozygous-floxed mutations for Dgrc8 (*Brn4-Cre/+; Dgrc8^flox/+^*) to obtain conditional knockout embryos (*Brn4-Cre/+; Dgrc8^flox/flox^*). *Brn4-Cre/+; Dgrc8^flox/+^, Dgrc8^flox/flox^* and *Dgrc8^flox/+^*embryos were considered as controls. The morning of the vaginal plug was considered as embryonic day (e)0.5. The embryos were collected at embryonic day e9.5, e11.5 and e13.5.

Genotyping of the *Dgcr8^flox^*, *Brn4-Cre* and *Rosa26-EYFP* alleles was performed using HighQu SampleIN Direct PCR Kit with the following primers: *Dgcr8^flox^-F* and *Dgcr8^flox^-R*, producing a 530-bp amplicon in the wildtype (WT), a 800-bp amplicon for the floxed allele and a 120-bp amplicon for the Cre-deleted Dgcr8 allele; *Brn4-Cre-F* and *Brn4-Cre-R* producing a 521-bp amplicon when Cre is present; and the *Rosa26-EYFP-1, Rosa26-EYFP-2* and *Rosa26-EYFP-3* producing a 600-bp amplicon for the WT allele and a 320-bp amplicon when the transgene is present (See Table S1 for primer sequences).

### Sample preparation scRNAseq analysis

A complete list of embryo genotypes used for scRNAseq is provided in Table S2. One set of e11.25 (1 control, 2 mutants) and e13.25 (2 controls, 2 mutants) embryos was obtained from timed-matings of *Dgcr8^flox/flox^* females with *Brn4-Cre; Dgrc8^+/flox^* males. Another set of e11.25 (2 controls, 1 mutant) and e13.25 (1 control, 2 mutants) embryos was obtained from timed matings of *Dgcr8^flox/flox^; Rosa26-YFP^tg/tg^* females with *Brn4-Cre; Dgrc8^+/flox^*males. Embryo genotypes were determined by presence of YFP fluorescence and PCR genotyping. Neural tube from selected embryos were dissected from the cervical to thoracic regions in Hibernate®-E medium (Thermofisher, A12476-01) supplemented with 0.5 mM GlutaMAX™-I (Thermofisher, 35050038) and 2% B-27® (Thermofisher, 17504001). Samples were incubated in FACSmax cell dissociation solution (Fisher Scientific, T200100) containing 10x Papain (30 U/mg, Sigma-Aldrich, 10108014001) for 15 min at 37°C to dissociate the tissue. Cells were subsequently transferred to resuspension buffer consisting of sterile 1× PBS (Gibco, 14190169) supplemented with 2% fetal bovine serum (Anprotec, AC-SM-0143) to obtain a single-cell suspension. Cell clumps were mechanically dissociated by gentle pipetting and filtered sequentially through 40 μm cell strainers (EASYStrainer™, Greiner, 542040) and 30 μm strainers (Miltenyi Biotec, 130-041-407). Sample quality was assessed by measuring cell viability and the presence of residual clumps. Only samples with viability greater than 80% were used for sequencing.

### Single cell sequencing

A suspension of 8,000 single cells was loaded onto the 10x Genomics Single Cell 3′ Chip, and cDNA synthesis and library construction were performed as per the manufacturer’s protocol for the Chromium Single Cell 3′ v3 protocol. 150 bp paired end sequencing using an Illumina NovaSeq 6000 and NovaSeq X Plus was performed by the NGS core unit at the Institute of Human Genetics of the FAU Erlangen-Nürnberg according to the manufacturer’s protocol.

Demultiplexing, alignment to transcriptome version mm10-2020-A, filtering, as well as barcode and UMI counting, were performed using Illumina Dragen (v.4) Software and Cellranger (version 7.2.0, 10x Genomics, under default settings). Single-cell RNA sequencing datasets were processed using the Seurat v5 workflow in R (see Supplemental Experimental Procedures). A metadata table containing embryo identity, genotype, experimental category, and sequencing run information was generated prior to analysis. Individual datasets were loaded as Seurat objects and subjected to quality control filtering. Cells with fewer than 2,500 or more than 20,000 detected genes, or with mitochondrial transcript content greater than 6%, were excluded. Putative doublets were identified and removed using DoubletFinder with default parameters.

Filtered datasets were integrated using canonical correlation analysis (CCA) implemented in Seurat v5 (*106*). Clustering and dimensionality reduction analyses were performed on the integrated dataset, and cluster identities were assigned based on differential gene expression analysis using Seurat’s FindAllMarkers function together with known marker gene expression visualized by DotPlot analysis. Iterative subclustering was subsequently performed on transcriptionally heterogeneous populations, including dI6/V1 interneurons, dI3 interneurons, and dorsal progenitor populations, to improve cell-type resolution and refine cluster annotations. Final cell identities were grouped into broader macro-categories corresponding to neuronal, progenitor, neurogenic, and non-neural populations. Complete R scripts detailing the performed analysis including the applied thresholds are available as Supplemental Experimental Procedures. Results of the DGE analysis in Figs. 3B,C,E and 4A are available in Data S1.

### Target prediction analysis

Target prediction analysis for miR-9 and let-7 was performed using publicly available TargetScan datasets for mouse (*78*). For interspecies comparisons, target predictions were obtained from the corresponding TargetScan datasets for human and fish. For frog and chicken, target predictions were based on the mouse datasets due to the absence of corresponding datasets in TargetScan. Predicted targets were subsequently used for comparative analyses across species (see Supplemental Experimental Procedures).

### Plasmids

The reporter and expression plasmids pCAG-nDsRedFL-intron and pCAG-nDsRedFL-miR9/9* were a gift from Rikako Sanuki (Addgene plasmids #177804 and 177805; RRIDs: Addgene_177804 and Addgene_177805) (*107*). The pCAG-nDsRedFL-miR9_sponge plasmid was generated by cloning two copies of the miR-9 sponge sequence from pBabe-puro-miR-9 sponge, which was a gift from Bob Weinberg (Addgene plasmid # 25040; RRID:Addgene_25040) (*108*), into the pCAG-nDsRedFL-intron backbone. The pCAG-nDsRedFL-let7c plasmid was generated by cloning the let-7c sequence into the pCAG-nDsRedFL-intron backbone. The pCI-Lin28a-IRES-H2B-GFP plasmid was generated by cloning the mouse Lin28a coding sequence into the pCI-H2B-GFP backbone, a gift from Tatjana Sauka-Spengler (Addgene plasmid #92399; RRID:Addgene_92399) (*109*).

For luciferase reporter assays, psiCHECK3-Unstructured (gift from Anthony Leung; Addgene #136025, RRID:Addgene_136025) (*110*) was used as control vector. For psiCHECK3-3′UTR-OC2.1, psiCHECK3-3′UTR-OC2.2, psiCHECK3-3′UTR-Pou2f1, psiCHECK3-3′UTR-Pou2f2.1, psiCHECK3-3′UTR-Pou2f2.2 and psiCHECK3-3′UTR-Zfhx3 fragments corresponding to the 3’-UTRs of these transcripts were cloned into psiCHECK3-Unstructured. The Pou2f1 3’-UTR fragment was synthesized as gBlocks fragment by IDT and contains all predicted miR-9 sites plus approximately 100bp of surrounding 5’ and 3’ sequence. The other fragments were cloned by amplifying the corresponding regions by PCR from genomic DNA. All plasmid constructs generated in this study were assembled using the NEBuilder HiFi DNA Assembly Cloning Kit according to the manufacturer’s instructions. Details about plasmids and primers used for cloning are available in Tables S1 and S3.

### Mouse embryonic stem cell culture

Mouse embryonic stem cell (mESCs) culture and differentiations were performed as previously described (*8*, *60*, *111*, *112*). HM1 cells were used for the experiments. 60mm x 15mm Corning Suspension culture dishes were precoated with 0.1% gelatine in DPBS. mESCs were maintained on mitotically inactivated mouse embryonic fibroblasts (feeders), supplemented with ES media and LIF (1000units/ml). For initiating the differentiation, mESCs were dissociated with 0.05% trypsin, followed by panning for 30 mins to remove the feeders. 60,000 to 80,000 cells were plated on 35mm x 10mm Corning cell bind dishes coated with 0.1% gelatine. Cells were supplemented with N2B27 media with 10ng/ml FGF2/bFGF. At day2 (D2), the media was changed to N2B27 supplemented with FGF2/bFGF (10ng/ml), CHIR99021 (5uM), DMH1 (2uM) and SB431542 (10uM). After D2 medium was changed daily. For generating neurons with dorsal identity, cells were kept in N2B27 with 100nM RA from D3-D6 and maintained in N2B27 until D13. For imparting ventral identity, cells were kept in 100 nM RA and 500nM SAG until D6 and N2B27 + 500nM SAG from D6-D10. From D10-D13 cells were maintained in N2B27.

### Reverse transcription and quantitative real-time PCR (RT-qPCR)

RT-qPCR was performed to quantify the expression of both mature miRNAs and pri-miRNA host transcripts. Total RNA was reverse transcribed using two different approaches depending on the target. For mature miRNAs, cDNA synthesis was performed using miRNA-specific TaqMan™ reverse transcription primers according to the manufacturer’s instructions. For pri-miRNA host transcripts, 2,000 ng of total RNA was reverse transcribed using random hexamer primers and the High-Capacity cDNA Reverse Transcription Kit (Applied Biosystems) according to the manufacturer’s instructions. Reverse transcription was carried out under the following conditions: 25°C for 10 min, 37°C for 120 min, 85°C for 5 min, followed by a 4°C hold.

Quantitative PCR was performed in 96-well plates using a total reaction volume of 10 μL containing 5 μL TaqMan™ Fast Advanced Master Mix, 0.5 μL TaqMan™ assay/probe, 1 μL cDNA, and 3.5 μL nuclease-free water. All reactions were performed in duplicate technical replicates. For mature miRNA quantification, fluorescence was detected using the FAM reporter dye. Cycling conditions consisted of an initial incubation at 50°C for 2 min, followed by denaturation at 95°C for 20 s and 40 cycles of 95°C for 3 s and 60°C for 30 s. For pri-miRNA host transcript quantification, fluorescence was detected using SYBR Green. Reactions were incubated at 50°C for 2 min, followed by initial denaturation at 95°C for 2 min and 40 amplification cycles of 95°C for 30 s and 60°C for 1 min. Relative expression levels were calculated using the 2^−ΔΔCt method. snoRNA was used as the endogenous control for mature miRNA analysis, whereas ACTB (β-actin) served as the housekeeping gene for pri-miRNA host transcript analysis. Details about material used for RT-qPCR are available in Table S3. RT-qPCR data underlying Fig. S7B-E is available in. Data S3.

### Luciferase assays

Luciferase assays were performed using a Dual-luciferase Reporter assay system on HEK293T cells. Cells were seeded at a density of 2×10^4^ cells per well in poly-L-lysine (0.1 mg/mL)-coated, clear-bottom 96-well plates. Twenty-four hours after seeding, cells were co-transfected with the pCAG-DsRed-miR-9/9* expression construct and psiCHECK-3 reporter plasmids containing the respective 3′UTRs using Lipofectamine™ 3000 (Thermo Fisher Scientific) according to the manufacturer’s instructions. Transfections were performed in Opti-MEM™ with a final volume of 100 μL per well, consisting of 90 μL Opti-MEM™ and 10 μL transfection mixture. Each condition was analysed in three technical replicates. Twenty-four hours after transfection, the medium was replaced with 100 μL of fresh DMEM. Luciferase activity was measured 48 hours post-transfection using Dual-Glo Luciferase Reporter Assay System (Promega, E2920) according to the manufacturer’s instructions. Relative luciferase activity was calculated as the ratio of Renilla to Firefly luciferase luminescence (Renilla/Firefly), with Firefly luciferase serving as the internal normalization control. Details about material used for Luciferase assays are available in Table S3. Data underlying graphs in Fig. S6B is available in Data S2.

### In-ovo electroporation

In-ovo electroporation was performed at Hamburger-Hamilton (HH) stage HH15-16 and embryos were collected at HH25 or HH27 (48h or 72h after electroporation). Electroporated plasmids were pCAG-nDsRedFL-intron, pCAG-nDsRedFL-miR9/9*, pCAG-nDsRed-let7c, pCI-Lin28b-H2B-GFP (1 µg/µL) and pCAG-nDsRed-mir9_sponge (1.5 µg/µL). The DsRed or GFP was used to visualize electroporated cells and assess the electroporation efficiency.

### Tissue collection

Collected embryos were fixed in ice-cold 4% paraformaldehyde (PFA) in PBS (overnight for RNAscope samples; 30 min, 2h30min or 3h (e9.5, e11.5, and e13.5, respectively) for mouse embryo immunostainings and 1h for chicken embryo immunostainings). Following fixation, mouse embryos were washed according to developmental stage. Embryos at E11.5 were washed twice for 5 min in 0.24M phosphate buffer (PB) followed by one 5-min wash in 0.12M PB. Embryos at E13.5 were washed for 5 min in PBS, followed by three consecutive 5-min washes each in 1M, 0.48M, and 0.24M PB, and finally two 5-min washes in 0.12M PB. Chicken embryos were washed three times in PBS. After washing, samples were cryoprotected overnight at 4°C in 15% sucrose prepared in 0.12M PB for mouse embryos and 30% sucrose in PBS for chicken embryos. For RNAscope, spinal cords were subsequently dissected in 15% sucrose/0.12M PB and allowed to sink completely before embedding in a 1:1 solution of OCT and 15% sucrose/0.12M PB. For immunostaining, embryos were embedded and frozen in 1M PB/7.5% gelatine/15% sucrose solution. Embedded samples were frozen, stored at −80°C until sectioning.

### Immunofluorescence labelling and RNAscope *in situ* hybridization (ISH)

Immunolabeling and in ISH were performed on 14 µm serial cryosections as previously described^9^. The primary and secondary antibodies used for immunolabelling are listed below (see Table S2). ISH was performed using the RNAscope Multiplex Fluorescent Reagent Kit v2 (ACDBio; #323270) and the RNAscope™ Plus smRNA-RNA HD Detection Kit (ACDBio; #322780) according to the manufacturer’s instructions, with minor modifications. Specifically, the protease treatment was adjusted according to the embryonic stage. Protease III was used for both e11.5 and e13.5 tissues with an incubation time of 3 min and 5 min for e11.5 and e13.5, respectively. Details about antibodies and RNA scope probes are available in the Supplemental Table S3.

For combined immunohistochemistry and RNAScope, immunohistochemistry was performed following RNAscope detection of miR-9 with Protease III treatment shortened to 1 minute at 40 degrees. After RNAscope, spinal cord sections were washed 5 times, 2 minutes each with RNAScope wash buffer and sections were incubated in co-detection blocker (ACD) for 30 minutes at room temperature, followed by incubation in 1% BSA in 1X PBS+ 0.1% Triton-X-100. Slides were incubated overnight at 4 degrees with Pou2f2, Onecut2, Zfhx3 or Neurod2 primary antibodies, diluted (1:200) in co-detection antibody diluent. Secondary antibodies in combination with DAPI were also diluted in co-detection antibody diluent (also 1:200) and slides were kept at room temperature for 2 hours. Finally, slides were washed 10 times, 5 minutes each, with 1X PBS+ 0.1 % Triton-X-100 before mounting and imaging.

### EdU birth-dating

EdU labelling was performed via intraperitoneal injection of 30 mg/kg body weight (resuspended in 1xPBS) into mice at E10.5 and e11.5. Embryos were collected afterwards at e13.5 for further analysis. EdU was detected using Alexa647 Click-iT EdU Imaging Kit (Invitrogen #C10340) according to the manufacturer’s specifications

### Imaging and quantifications

Immunohistochemistry images were acquired using Zeiss Observer microscope equipped with an Apotome.2 structured illumination module and a 20× air objective [numerical aperture = 0.8].

### Quantification of total number of cells and mean cells

Nuclei were segmented with StarDist and manually corrected in Fiji. Individual channel intensities were extracted using the ROI Manager’s Multi Measure tool in Fiji and exported as CSV or Microsoft Excel files. Data was analyzed and plotted using R (version 4.5.2). Within each section, nuclear intensities were normalized to a scale of 0 to 1, setting the lower and upper bounds after removing a defined quantile of the dimmest and brightest signals for image normalization (see Table S4). Objects were counted as positive for expression of temporal TFs if their normalized intensities exceeded a normalized intensity threshold (see Table S4). Plots were generated using ggplot2 (version 3.4.2).

To quantify miR-9 intensity density distributions of areas positive for the other respective markers areas outside the spinal cord grey matter were removed from the analysis and a 4000×4000 pixel image encompassing the entire spinal cord was down-sampled to 200×200 pixels with bilinear interpolation in Fiji. Intensities for miR-9 and the other markers in the down-sampled image were afterwards normalized between 0 and 1, as described above. tTFs were afterwards partitioned into positive and negative areas based on a normalized intensity threshold of 0.2, and density distributions of miR-9 intensities in the positive areas were plotted using ggplot’s function geom_freqpoly with bins = 30.

Quantification of number of pHH3 and cC3 was performed using the Cell Counting tool in Fiji/ImageJ2 v2.10.0, and plots were generated using the Prism Software (version 10.6.0).

### Area measurements

Spinal cord areas were quantified by manually defining tissue boundaries as ROIs using the Freehand selection tool in Fiji/ImageJ2 v2.10.0. The neuronal area was subtracted from both the total area and the progenitor area. Raw area values (µm^2^) were exported as Microsoft Excel files and subsequently analyzed to calculate the corresponding percentages. Plots were generated using the Prism Software (version 10.6.0).

### Experimental design and statistical analysis

Experiments were not randomized, and no statistical method was used to predetermine sample size. The investigators were not blinded during experiments and outcome assessment. For quantifications of *in ovo* electroporation experiments, quantifications on the “electroporated” (targeted) side of the spinal cord were compared to quantifications on the “nonelectroporated” (contralateral) side of the same embryo. Quantifications in mouse embryonic models were performed at the cervical level of the spinal cord and cells were quantified on both sides of the spinal cord. A complete list of analyzed sections and embryos per genotype is provided in Table S4. Statistical analyses and plotting were performed in R (https://www.r-project.org) or Graphpad Prism (version 10.6.0). Applied statistical tests and p-values are included in figure panels and legends.

### Statement on the usage of artificial intelligence

During preparation of this work, the authors used the services of ChatGPT 5 and Google Gemini to assist with the writing R code for quantitative image and scRNAseq data analysis. The authors moreover used ChatGPT 5 to improve readability and clarify phrasing during the preparation of the manuscript. After using artificial intelligence interfaces, the authors reviewed and edited the content as needed and take full responsibility for the content of the publication.

## Acknowledgments

We thank J. Briscoe, R. B. Roome, J. Delas and E. Parasyraki for comments on the manuscript, J. Przibylla, Emily Brockmann and Arif Ekici for technical support and E. Sock, C. Lie, R. Beckervordersandforth, P. Soba, T. Müller, Susan Brenner-Morton and J. Briscoe for sharing mouse lines, lab reagents, antibodies, and equipment access. We are also grateful to all members of the Sagner lab, past and present, for help, advice, and critical feedback. We further acknowledge the services of the Targetscan database and Addgene.

## Funding

This work was funded by the: German Research Foundation (DFG) Emmy Noether Programme 455354162 (A.S.), German Research Foundation (DFG) Walter-Benjamin Programme (Project number: 573237392) (E.R.), Interdisciplinary Centre for Clinical Research IZKF - E36 and IZKF-S1 (both to A.S.) WBI.WORLD fellowship (E.R.).

## Author contributions

Conceptualization: E.R., L.C.S., A.S.M., A.S.

Methodology: E.R., A.S.M., M.P.,

Validation: E.R., L.C.S., A.S.M., M.P.,

Formal analysis: E.R., L.C.S., A.S.M., M.P., J.S., A.S.

Investigation: E.R., L.C.S., A.S.M., M.P., J.S.

Resources: H.-M. J., J.W., M.W.

Data curation: E.R., L.C.S., A.S.M., M.P., A.S.

Visualization: E.R., L.C.S., A.S.M., M.P., J.S., A.S.

Writing - original draft: E.R., A.S.

Supervision: A.S.

Project administration: A.S.

Funding acquisition: E.R., A.S.

## Competing interests

The authors declare they have no competing interests.

## Notes

### Competing Interest Statement

The authors have declared no competing interest.

