## Supplemental Figures and Tables for "microRNAs control temporal patterning of the vertebrate nervous system"

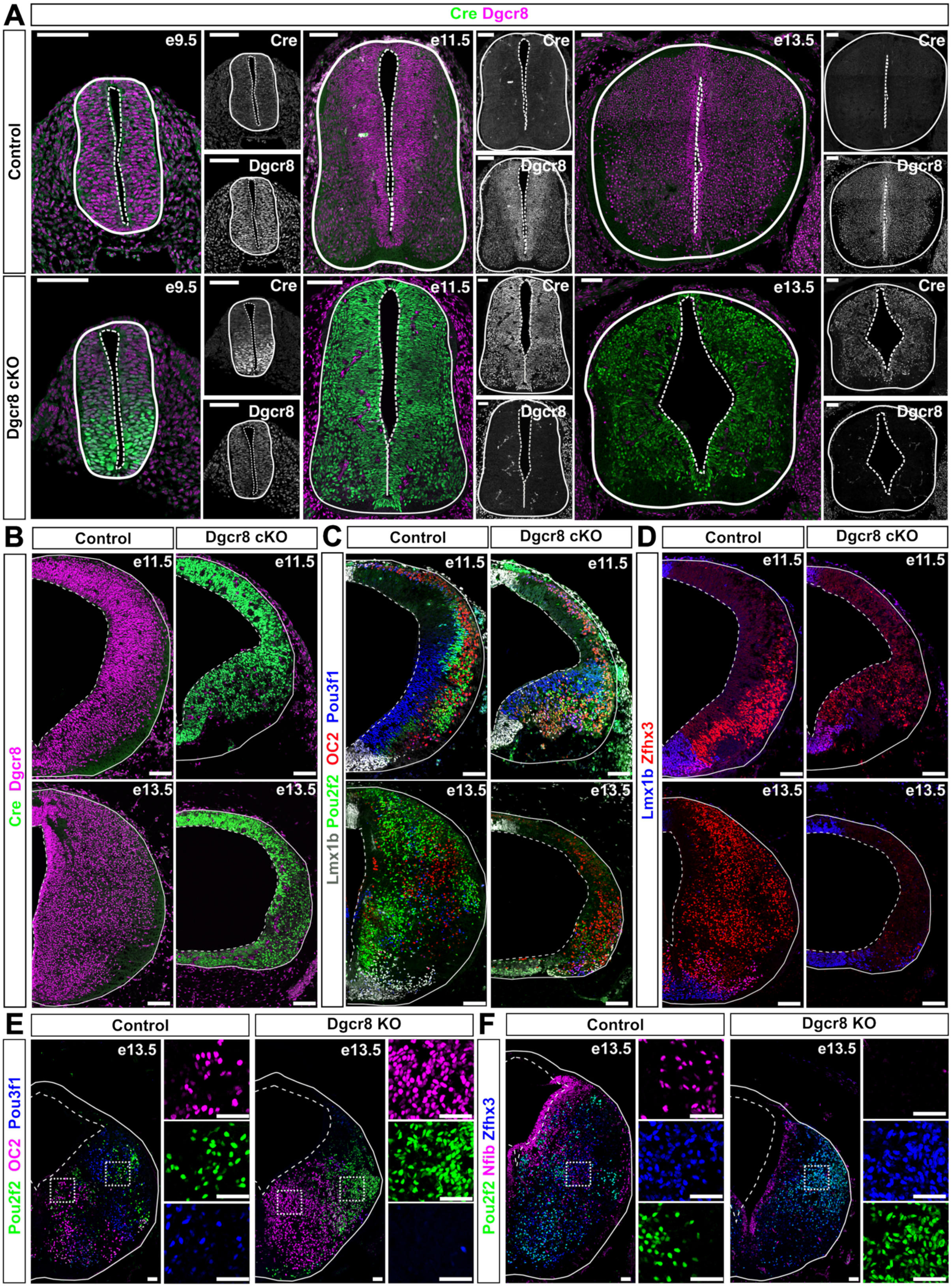

A
Cre Dgcr8
Control
Dgcr8 cKO
e9.5
e11.5
e13.5
Cre
Dgcr8
B
Control
Dgcr8 cKO
Cre Dgcr8
C
Lmx1b Pou2f2 OC2 Pou3f1
D
Lmx1b Zfhx3
E
Control
Dgcr8 KO
Pou2f2 OC2 Pou3f1
F
Pou2f2 Nfib Zfhx3

**Fig. S1. Further characterization of Dgcr8 cKO embryos (related to Fig. 1)**

**(A)** Dgcr8 is present throughout the spinal cord in control embryos (genotype Dgcr8 flox/flox). In Dgcr8 cKO embryos, Cre is present by e9.5 in the ventral spinal cord, but Dgcr8 is still detectable at this stage. At e11.5, Cre is present and Dgcr8 undetectable throughout the spinal cord. **(B-D)** Characterization of midbrain development in Dgcr8 cKO embryos. **(B)** Staining of control (left) and Dgcr8 cKO embryos (right) at e11.5 (top) and e13.5 (bottom). Dgcr8 is present throughout the midbrain in control and undetectable in the Dgcr8 cKO embryos. Also note the strong reduction in overall nuclei number in the Dgcr8 cKO embryos. **(C)** Staining for the dopaminergic neuron marker Lmx1b and the early tTFs OC2, Pou2f2 and Pou3f1. The fraction of neurons expressing OC2 is elevated in Dgcr8 cKO embryos. **(D)** Staining of embryonic midbrains for Lmx1b and Zfhx3. **(E, F)** Characterization of hindbrain development in Dgcr8 cKO embryos. **(E)** e13.5 hindbrain section of control (left) and Dgcr8 cKO embryos (right) stained for OC2, Pou2f2 and Pou3f1. Similar to the spinal cord, Pou3f1 is almost completely absent, while OC2 and Pou2f2 expression are expanded. **(F)** Staining of e13.5 hindbrain sections for Pou2f2, Zfhx3 and Nfib. Nfib levels are reduced in the hindbrains of Dgcr8 cKO embryos. Scale bars = 100 μm,

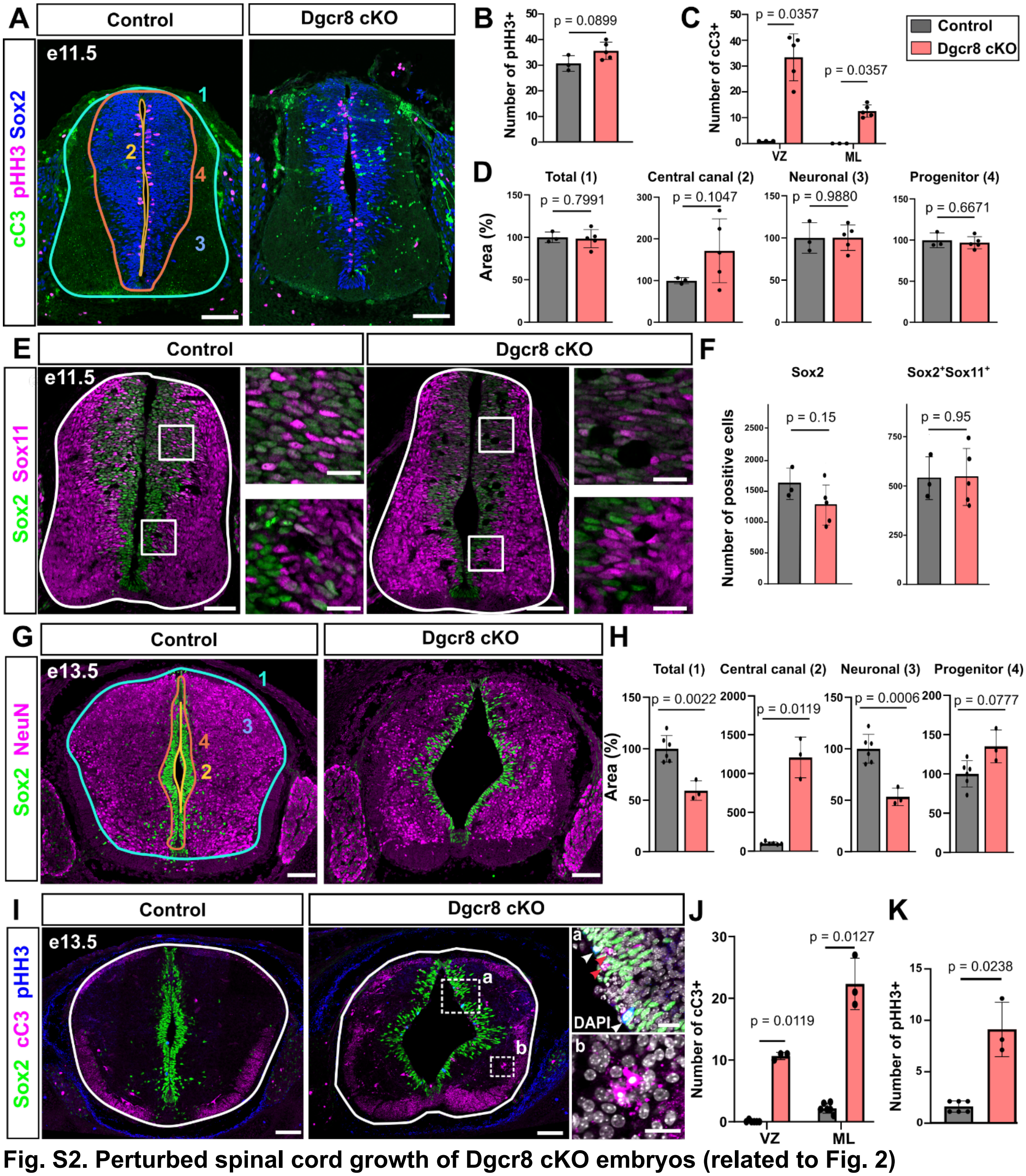

**Fig. S2. Perturbed spinal cord growth of Dgcr8 cKO embryos (related to Fig. 2)**

**(A)** e11.5 cervical spinal cord section of control (left) and Dgcr8 cKO (right) embryos stained for the progenitor marker Sox2, apoptotic marker cleaved Caspase 3 (cC3) and proliferation marker phospho-Histone H3 (pHH3). **(B, C)** Quantification of proliferative pHH3+ and apoptotic cC3+ cells. Apoptotic cells were separately quantified in the ventricular zone (VZ) and mantle layer (ML). **(D)** Neither total area nor the area of the VZ and ML are altered in Dgcr8 cKO embryos at e11.5. The size of the individual compartments was normalized to the respective average size of the compartment in control embryos. **(E)** e13.5 cervical spinal cord sections of control (left) and Dgcr8 cKO (right) embryos stained for progenitor marker Sox2 and neuronal marker NeuN. **(F)**

Total area, as well as area of the central canal and ML are altered in Dgcr8 cKO embryos. VZ area is not significantly altered in the Dgcr8 cKO embryos. We note however, that the VZ often appears fragmented. **(G)** e13.5 cervical spinal cord sections of control (left) and Dgcr8 cKO (right) embryos stained for Sox2, cC3, pHH3. **(H, I)** Increase in the number of apoptotic cC3+ cells (H) and proliferating pHH3+ cells in Dgcr8 cKO embryos at e13.5. White arrowheads point Sox2+pHH3+ cells, while red arrowheads point to Sox2-cC3+ cells in the VZ.
Scale bars = 100 µm, insets = 25 µm, Statistical significance was assessed by Student's t-test and Wilcoxon rank test, p-values are integrated on the graphs.

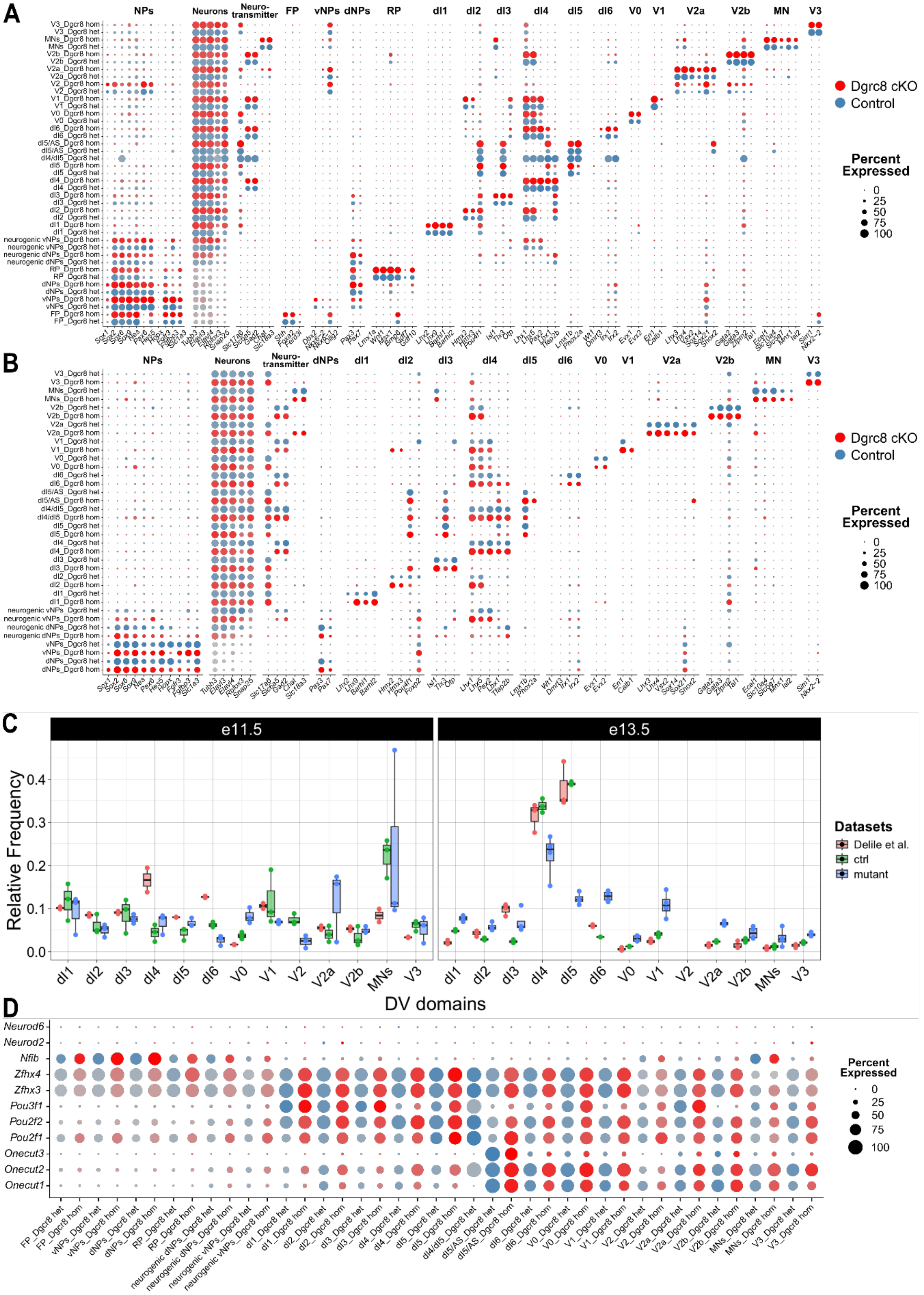

**Fig. S3. Spatial patterning is not strongly perturbed in Dgcr8 cKO embryos (related to Fig. 3)**

**(A, B)** Dotplots indicating expression of progenitor and neuronal markers, neurotransmitters and markers for the different DV classes of spinal cord neurons in control (blue) and Dgcr8 cKO (red) in the e11.25 (A) and e13.25 (B) scRNAseq data. **(C)** Relative frequency of neurons belonging to different DV classes in scRNAseq data from Delile et al. ([10]), and in the datasets from control and Dgcr8 cKO embryos at e11.25 and e13.25. Relative frequencies in our e11.25 and e13.25 datasets were compared frequencies in the e11.5 and e13.5 datasets from Delile et al. ([10]), respectively. **(D)** Dotplot of early tTFs indicating comparable expression in different spinal cell populations in control and Dgcr8 cKO embryos at e11.25.

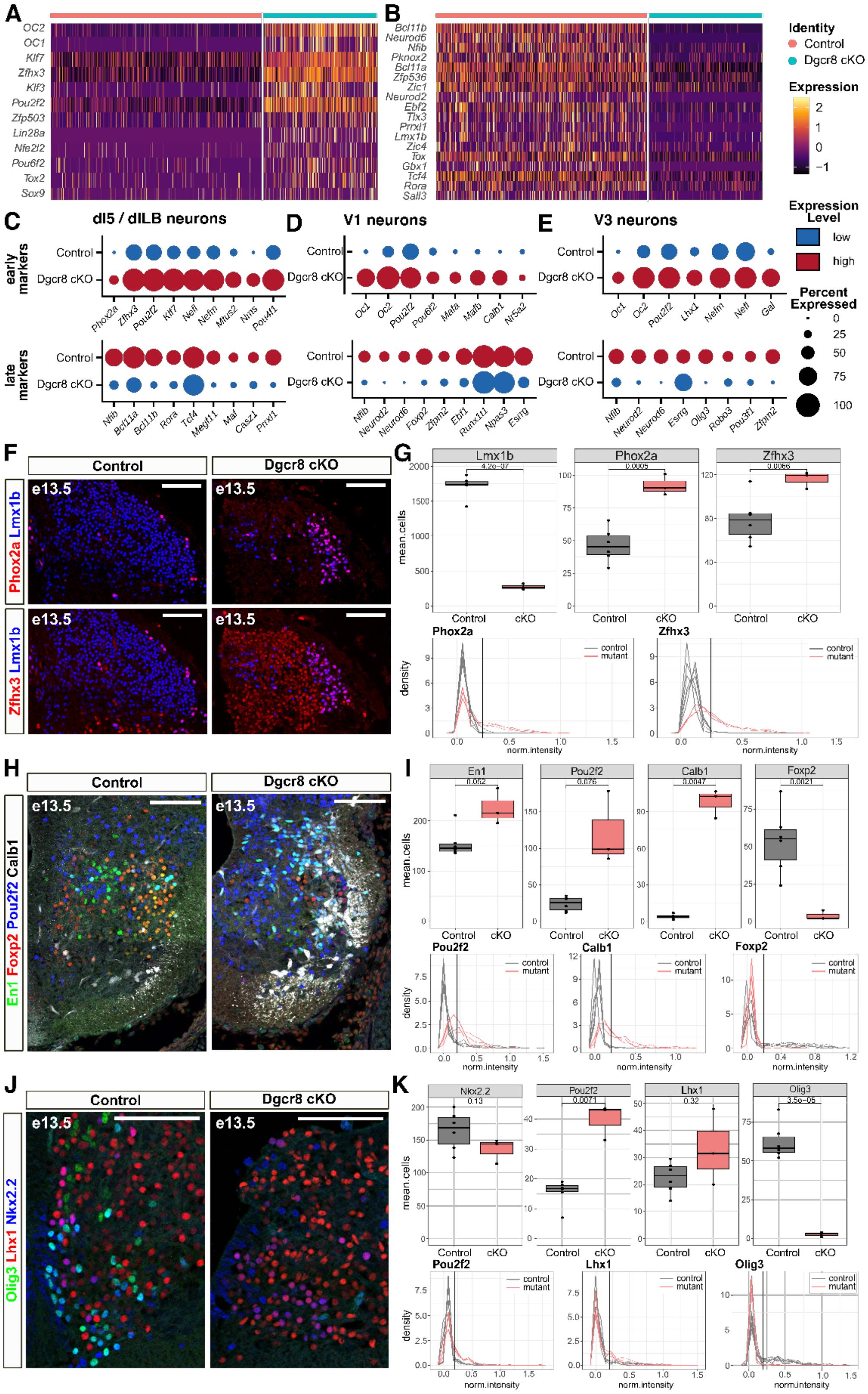
A
B
Identity
Control
Dgcr8 cKO
Expression
C
dI5 / dILB neurons
D
V1 neurons
E
V3 neurons
Expression Level
low
high
Percent Expressed
early markers
late markers
F
Control
Dgcr8 cKO
e13.5
Phox2a Lmx1b
Zfhx3 Lmx1b
G
Lmx1b
Phox2a
Zfhx3
mean.cells
density
norm.intensity
control
mutant
H
En1 Foxp2 Pou2f2 Calb1
I
En1
Pou2f2
Calb1
Foxp2
J
Olig3 Lhx1 Nkx2.2
K
Nkx2.2
Pou2f2
Lhx1
Olig3

**Fig. S4. Temporal shifts in neuronal subtype specification in Dgcr8 cKO embryos (related to Fig. 3)**

**(A, B)** Heatmaps indicating expression levels of up- (A) and down-regulated TFs (B) in neurons from Dgcr8 cKO embryos in the e13.25 scRNAseq data **(C-E)** Dotplots indicating expression of early (top) and late (bottom) neuronal subtype markers in dI5/dILB (C), V1 (D) and V3 (E). (**F, H, J)** e13.5 cervical spinal cord sections stained for DV class (Lmx1b [F], En1 [H], and Nkx2.2 [J]) and temporally specified neuronal subtype markers (Phox2a and Zfhx3 [F], Calb1, Pou2f2 and Foxp2 [H], and Lhx1, Pou2f2 and Olig3 [J]). **(G, I, K)** Quantification of the number of neurons expressing the respective markers (top) and density plots showing the intensity distributions for the indicated markers in the respective neuronal populations. The black line on the intensity distributions indicates the threshold above which neurons were counted as positive.
Scale bars in F, H, J = 100 µm, statistical significance in G, I, K was assessed by unpaired t-tests, p-values are integrated on the graphs.

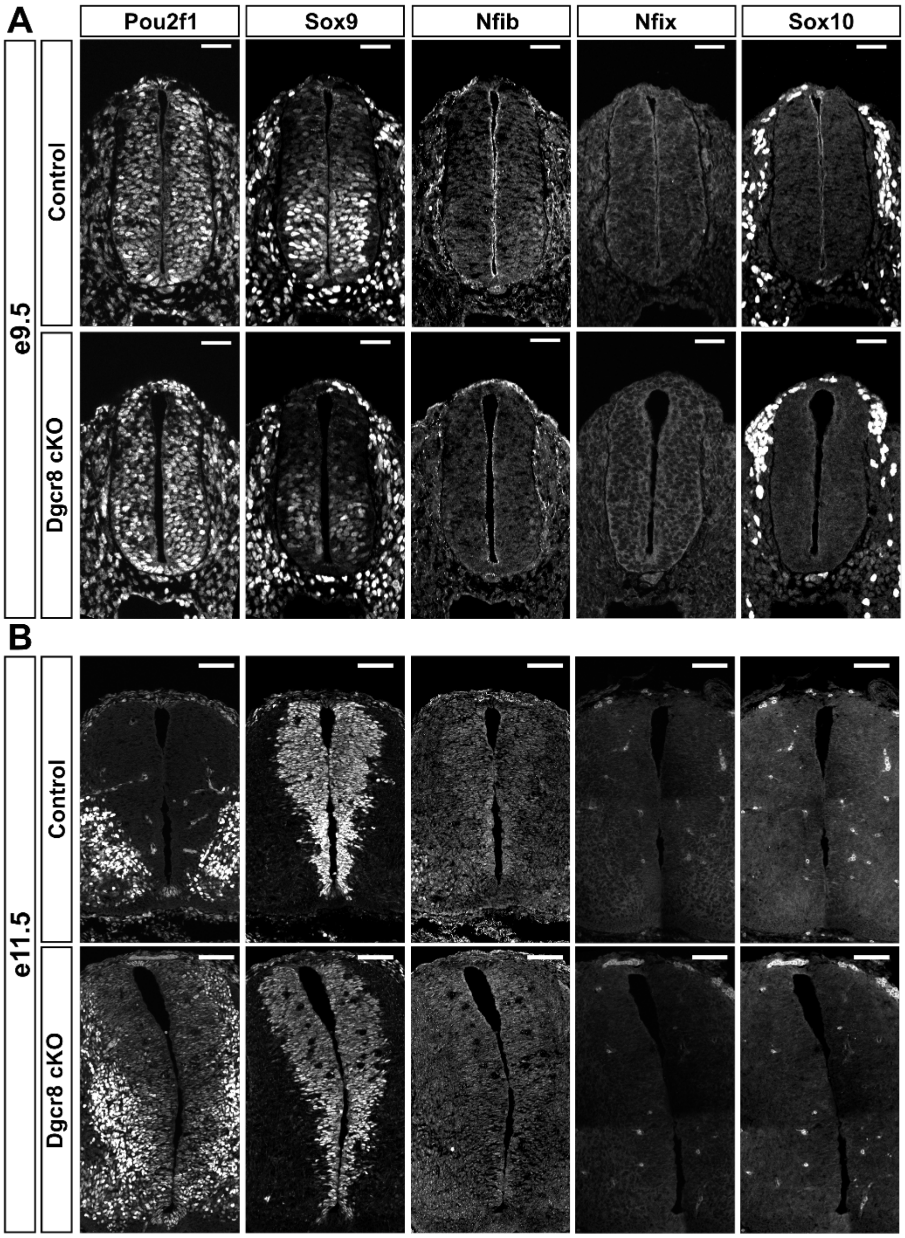

**Fig. S5. Sequential induction of the gliogenic program is perturbed in Dgcr8 cKO embryos (related to Fig. 4)**

**(A, B)** e9.5 (A) and e11.5 (B) cervical spinal cord sections of control (top) and Dgcr8 cKO (bottom) stained for the early progenitor marker Pou2f1 and the late progenitor markers Sox9, Nfib, Nfix and OPC marker Sox10. Scale bars = 100 µm

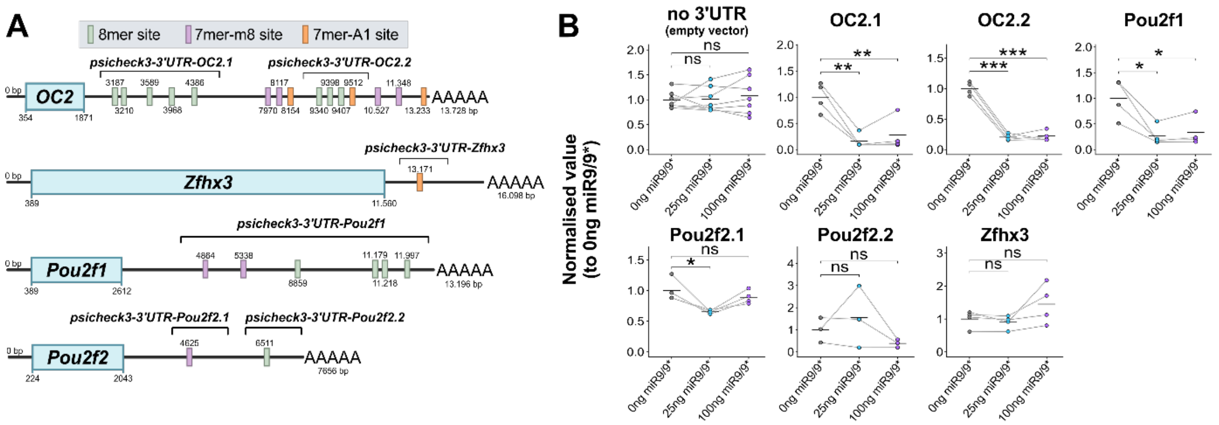

**C**

**Human**

| mir-9-5p Target gene | Conserved sites | Poorly conserved sites | let-7-5p Target gene | Conserved sites | Poorly conserved sites |
|---|---|---|---|---|---|
| OC2 | 13 | 7 | HMGA2 | 7 | 0 |
| YBX3 | 2 | 0 | ARID3B | 5 | 0 |
| LYVE1 | 1 | 3 | LIN28B | 4 | 1 |
| POU2F1 | 6 | 3 | FIGN | 6 | 0 |
| OC1 | 13 | 3 | TRIM71 | 7 | 1 |
| SLC50A1 | 2 | 0 | NR6A1 | 4 | 1 |
| POU6F2 | 4 | 0 | THRSP | 1 | 1 |
| TRPM7 | 2 | 0 | USP44 | 3 | 0 |
| AP1S2 | 2 | 0 | FAM222B | 2 | 0 |
| PDK4 | 1 | 0 | IGDCC3 | 3 | 1 |
| LDLRAP1 | 3 | 0 | IGF2BP1 | 5 | 0 |
| CTNNA1 | 1 | 1 | VSTM5 | 2 | 0 |
| C2ORF15 | 1 | 1 | FZD3 | 2 | 1 |
| KCNJ2 | 2 | 0 | PXT1 | 1 | 1 |
| SGMS2 | 3 | 0 | DTX2 | 2 | 0 |

**Chicken**

| mir-9-5p Target gene | Conserved sites | Poorly conserved sites | let-7-5p Target gene | Conserved sites | Poorly conserved sites |
|---|---|---|---|---|---|
| OC2 | 12 | 1 | Hmga2 | 7 | 0 |
| OC3 | 3 | 0 | Trim71 | 3 | 0 |
| OC1 | 10 | 0 | Fign | 6 | 0 |
| Prmd6 | 2 | 0 | Lin28b | 5 | 0 |
| Ybx3 | 1 | 0 | OC3 | 2 | 1 |
| Prtg | 5 | 1 | Igf2bp3 | 2 | 2 |
| Mesdc1 | 1 | 0 | Bzw1 | 2 | 0 |
| POU2F1 | 6 | 0 | Bach1 | 2 | 0 |
| Wasf2 | 1 | 0 | Vstm5 | 1 | 0 |
| Ctnna1 | 1 | 0 | Yod1 | 1 | 0 |
| Ank2 | 1 | 0 | Prtg | 2 | 1 |
| Ccdc43 | 1 | 0 | Nr6a1 | 2 | 0 |
| Fbn1 | 2 | 0 | Adrb1 | 1 | 1 |
| Rab11a | 1 | 0 | Sall4 | 1 | 0 |
| Kcnmb2 | 1 | 1 | Fignl2 | 2 | 0 |

**Frog**

| mir-9-5p Target gene | Conserved sites | Poorly conserved sites | let-7-5p Target gene | Conserved sites | Poorly conserved sites |
|---|---|---|---|---|---|
| OC2 | 9 | 2 | Fignl2 | 2 | 0 |
| OC1 | 9 | 1 | Hmga2 | 3 | 0 |
| Prdm6 | 2 | 0 | Lin28b | 3 | 0 |
| OC3 | 1 | 0 | Prtg | 2 | 0 |
| Maea | 1 | 0 | OC3 | 2 | 0 |
| POU2F1 | 3 | 1 | Bzw1 | 2 | 0 |
| 4632415L05 Rik | 1 | 0 | Igf2bp3 | 2 | 2 |
| Kcnmb2 | 1 | 0 | Yod1 | 1 | 0 |
| Lamp3 | 1 | 0 | Skil | 2 | 0 |
| Fam19a5 | 1 | 0 | Pbx3 | 2 | 0 |
| Bhlhe41 | 1 | 1 | Igf1r | 2 | 0 |
| Foxp2 | 2 | 1 | Igdcc3 | 1 | 0 |
| Erg | 1 | 0 | Edn1 | 1 | 0 |
| Cpeb2 | 2 | 0 | Foxp2 | 3 | 0 |
| Rnf111 | 1 | 0 | Col4a6 | 1 | 0 |

**Fish**

| mir-9 Target gene | All sites | let-7 Target gene | All sites |
|---|---|---|---|
| onecutl | 5 | trim71 | 8 |
| osr1 | 5 | igdcc3 | 6 |
| onecut1 | 12 | si:dkey-157g16.6 | 3 |
| si:busm1-160c18.1 | 3 | arid3b | 6 |
| tgfbr2 | 2 | CABZ010 54965.1 | 3 |
| snai1a | 3 | fign | 4 |
| FRRS1 (1 of 3) | 2 | si:ch211-202m22.1 | 2 |
| itga5 | 5 | SEMA4A (2 of 2) | 5 |
| KCTD17 | 3 | slco1c1 | 2 |
| ARID3C | 4 | AL929007.1 | 3 |
| zgc:175175 | 2 | plagx | 3 |
| zgc:162144 | 3 | si:ch211-147h1.4 | 3 |
| b3gat3 | 2 | SMUG1 | 2 |
| CNNM2 (1 of 2) | 2 | NTN5 | 2 |
| ptpn21 | 3 | si:dkey-80c24.5 | 2 |

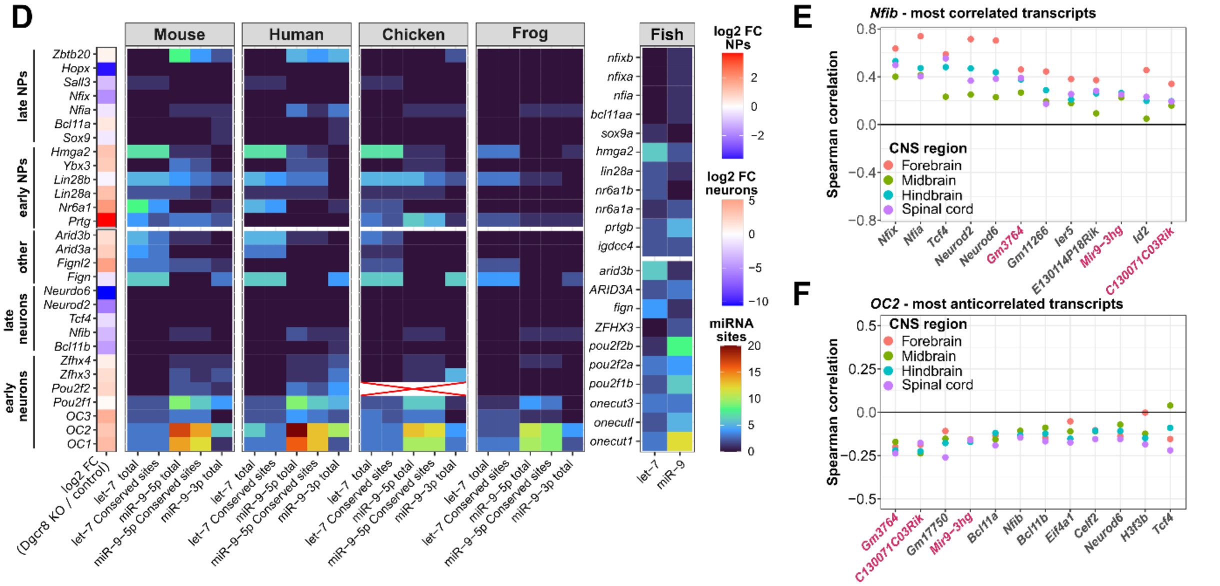

**Fig. S6. Evolutionary conserved antagonism of let-7 and miR-9 and the early transcriptional programs of progenitors and neurons (related to Fig. 5)**

**(A)** Schematics depicting the type and position of evolutionary conserved miR-9 sites in the 3'-UTRs of OC2, Pou2f1, Pou2f2 and Zfhx3 and the position and lengths of 3'-UTR fragments used for dual-luciferase assays (see B). Position of miR-9 sites and lengths of 3'-UTRs were obtained from annotations provided by Targetscan. **(B)** Relative repression by early tTF 3'-UTR fragments in response to 25 and 100 ng miR-9 expression plasmid measured in dual-luciferase assays. **(C)** Top 15 let-7-5p and miR-9-5p targets in human, chicken, frog and zebrafish (from left to right). Genes upregulated in the scRNAseq data from Dgcr8 cKO embryos are indicated by colored font. **(D)** Heatmap indicating Targetscan's Cumulative weighted context ++ score, which estimates the expected repression of an mRNA by a miRNA, for mRNAs encoding early and late progenitor and neuronal markers. A more negative cumulative weighted context ++ score indicates stronger expected repression. Statistical significance in B was assessed by paired t-tests. * < 0.05, ** < 0.01, *** < 0.001. **(E-F)** The miR-9 host genes *Gm3764*, *C130071C03Rik* and *Mir9-3hg* are amongst the genes, whose expression in neurons throughout the developing CNS correlates best with the late temporal marker *Nfib* (E) and is most anti-correlated with the early temporal marker *OC2* (F).

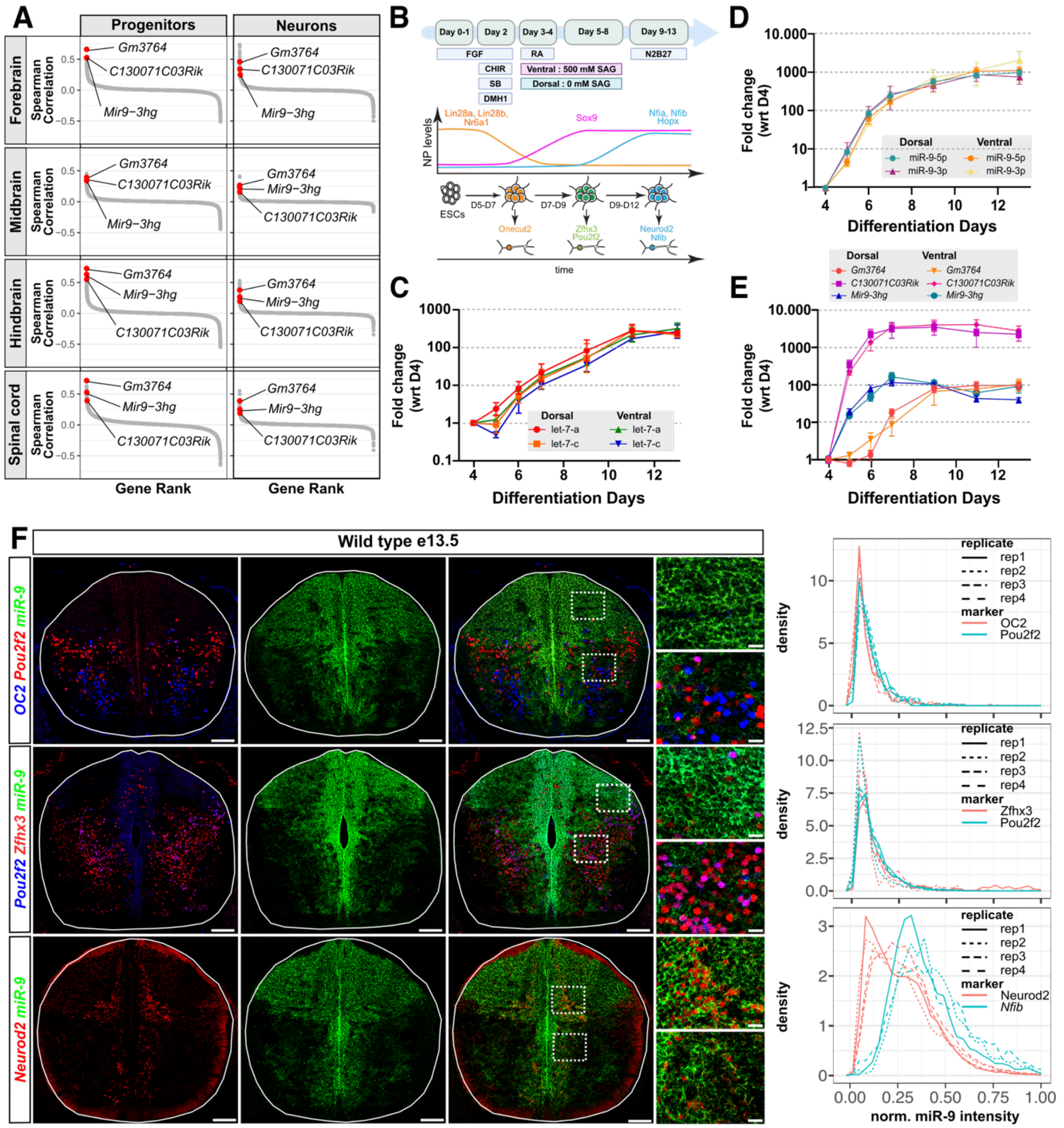

**Fig. S7. Further characterization of let-7 and miR-9 expression dynamics (related to Fig. 6)**

**(A)** Correlation rank plots depicting correlation between expression of the 3 miR-9 host genes *Gm3764*, *C130071C03Rik* and *Mir9-3hg* and *Nfib* in scRNAseq data from embryonic progenitors and neurons from the forebrain, midbrain, hindbrain and spinal cord. **(B)** mESC-based differentiation protocol for the generation of dorsal and ventral neural progenitors and neurons with a spinal cord identity (A). Scheme indicates expression dynamics of progenitor markers and chronological neuronal subtype specification in the in-vitro differentiations. Scheme drawn after Sagner et al. 2021 and Zhang et al. 2025 ([8,56]). **(C-E)** Expression dynamics of let-7 (C), miR-9-5p and miR-9-3p (D) and the mir-9 pri-miRNAs (E) in dorsal and ventral spinal cord differentiations. Data is depicted as relative fold change with respect to D4. Error bars represent standard error of

the mean. **(F)** miR-9 RNAscope in combination with immunofluorescence for OC2 and Pou2f2 (top row), Pou2f2 and Zfhx3 (middle panel) and Neurod2 (bottom panel). Plots on the right show normalized miR-9 intensity density distributions for areas positive for the indicated markers from spinal cord sections of 4 control embryos. miR-9 intensity distributions in *Nfib*+ areas were obtained by quantifying *Nfib* RNAscope intensities in spinal cord section from 3 control embryos (see Fig. 6D, top row)
Scale bars = 100 μm, insets = 25μm.

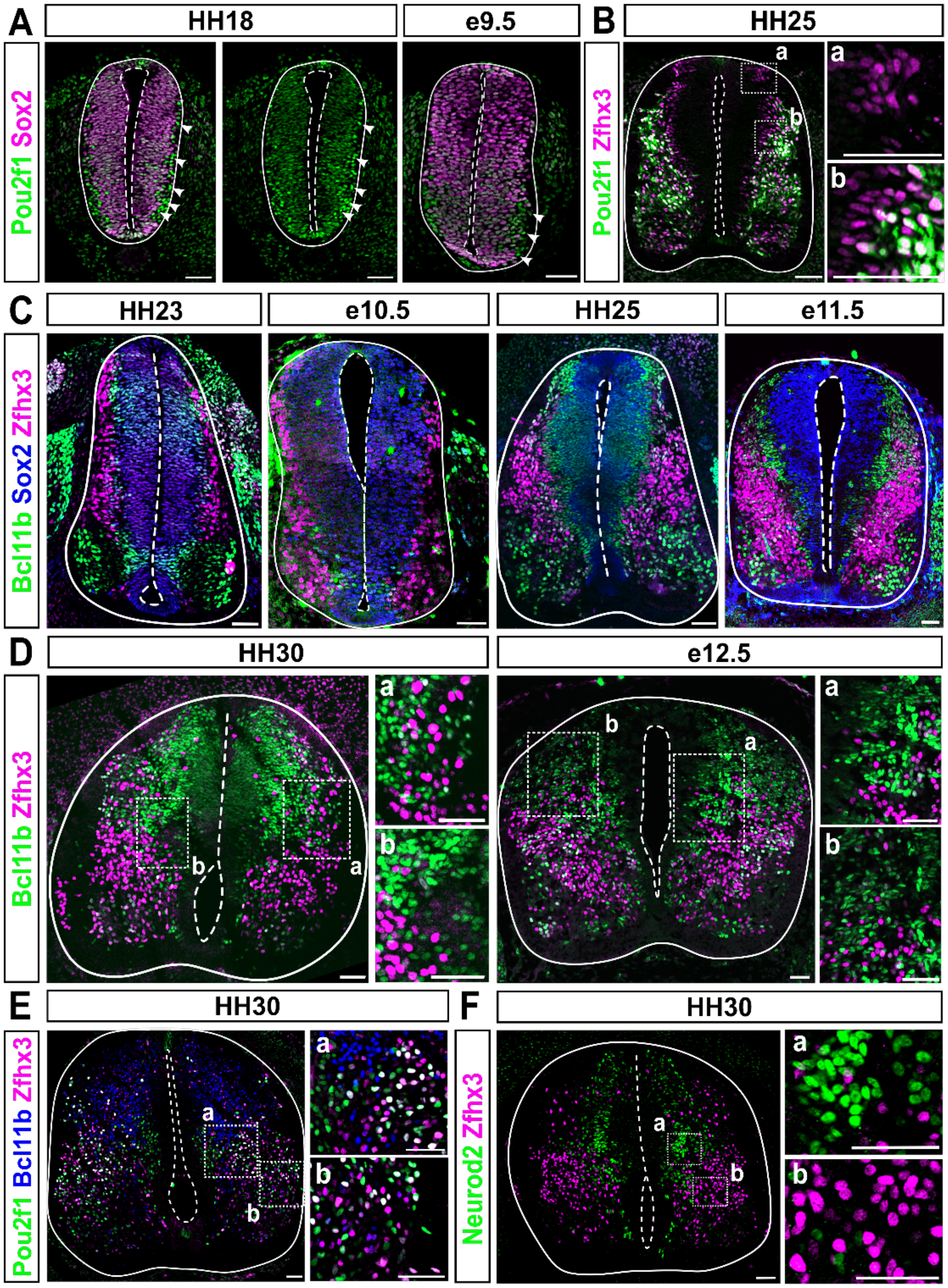

**Fig. S8. Conservation of temporal patterning in the embryonic chicken spinal cord (related to Fig. 7)**

**(A)** Chicken and mouse spinal cords at equivalent developmental stages (HH18 and e9.5) stained for Sox2, labelling the progenitors, and Pou2f1, labelling both progenitors (low levels) and neurons (high levels, arrowheads). Note similar distribution and mutually exclusive expression of both markers in neurons in both species. **(B)** Chicken spinal cord at HH25 stained for Pou2f1 and Zfhx3, labelling early- and intermediate-born neurons. **(C)** Chicken and mouse spinal cords at equivalent developmental stages (HH23 and e10.5 left, HH25 and e11.5 right) stained for the progenitor marker Sox2, the intermediate tTF Zfhx3 and Bcl11b, labelling late-born neurons. Note similar distribution and mutually exclusive expression of both markers in neurons in both species.

**(D)** Chicken and mouse spinal cord at equivalent developmental stages (HH30 and e12.5) stained for the intermediate tTF Zfhx3 and the late tTF Bcl11b. Note similar distribution and mutually exclusive expression of both markers in neurons in both species. **(E, F)** Mutually exclusive expression of early (Pou2f1 and Zfhx3) and late (Bcl11b and Neurod2) neuronal tTFs revealed by staining of chick embryonic spinal cords at HH30 for Pou2f1, Zfhx3 and Bcl11b (E), and Neurod2 and Zfhx3 (F). Scale bars = 50 μm

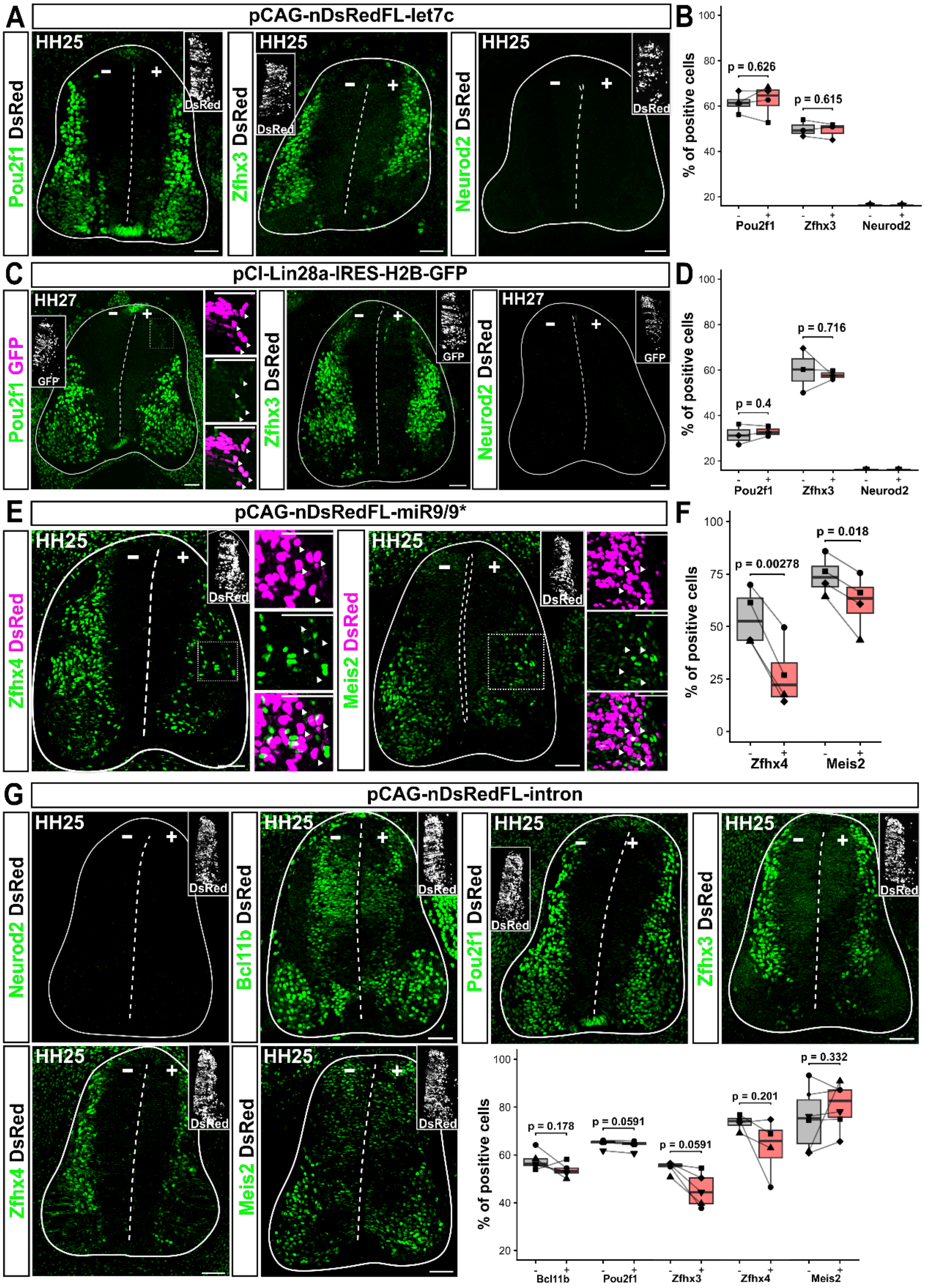

**Fig. S9. Perturbing let-7 expression or electroporation of a DsRed expression plasmid without miRNA hairpin do not perturb temporal patterning (related to Fig. 7)**

**(A, B)** Ectopic let-7c expression, indicated by presence of nuclear DsRed (nDsRed) does not repress early or induce late neuronal tTFs. **(C, D)** Interfering with let-7 biogenesis by ectopic expression of Lin28a does not repress early or induce late neuronal tTFs. We note however that Lin28a expression caused ectopic faint Pou2f1 expression in dorsal neurons in some sections (see left panel). **(E, F)** Ectopic miR-9 expression represses the early neuronal tTFs Zfhx4 and Meis2. **(G)** Electroporation of the nDsRed expression plasmid without miRNA hairpin does not perturb expression of Neurod2, Bcl11b, Pou2f1, Zfhx3, Zfhx4 and Meis2 expression. Scale bars = 50 μm; Statistical significance was assessed by paired t-test (B, D, F, G for Bcl11b, Meis2 and Zfhx3) or Wilcoxon signed rank test (G for Pou2f1 and Zfhx3).

**Table S1.**

List of primers used for genotyping and cloning.

| Primer name | Sequence | Purpose |
|---|---|---|
| Dgcr8$^{flox}$-F | GATCTCAGTAGAAAGTTTGGCTAAC | Genotyping of the Dgrc8 allele |
| Dgcr8$^{flox}$-R | GATATGTCTAGCACCAAAGAACTCC | Genotyping of the Dgrc8 allele |
| Brn4-Cre-F | CGA CCT GAA ACT GAA GTT AGC CTC C | Genotyping of the Brn4-Cre allele |
| Brn4-Cre-R | ATG CTCAGA AAA CGC CTG GCG ATC C | Genotyping of the Brn4-Cre allele |
| Rosa26-EYFP-1 | GGA GCG GGA GAAATG GAT ATG | Genotyping of the Rosa26-EYFP allele |
| Rosa26-EYFP-2 | AAA GTC GCT CTG AGT TGT TAT | Genotyping of the Rosa26-EYFP allele |
| Rosa26-EYFP-3 | AAG ACC GCG AAG AGT TTG TC | Genotyping of the Rosa26-EYFP allele |
| pCAG-nDsRedFL-let7c-F | cggtgatgcctttgagggatTAAGCTACAACTGAAGTTG | Cloning of let7c in pCAG-nDsRedFL |
| pCAG-nDsRedFL-let7c-R | cagatggacgcggccacgatACAAGCAGAATCCTAACC | Cloning of let7c in pCAG-nDsRedFL |
| DsRed-mir9 sponge_F | agacgatgacgataagtaactgtgtaggcggccgcctc | DsRed-mir9sponge cloning-step1 |
| mir9-Sponge x2_R | agagggaaaaagatctacactcgaggctgatcagcggg | DsRed-mir9sponge cloning-step1 |
| mir9spongeF-step2 | gctgatcagcctcgagtgtatgtgtaggcggccgcctc | DsRed-mir9sponge cloning-step2 |
| mir9spongeR-step2 | atttttggcagagggaaaaagatcttcgaggctgatcagcggg | DsRed-mir9sponge cloning-step2 |
| pCI-Lin28a-H2B-GFP_F | agctcatcgataagcttgatcagcagacgaccatgggc | Cloning Lin28a in pCI-H2B-GFP |
| pCI-Lin28a-H2B-GFP_R | ccgggctgcaggaattcgattcaattctgggcttctggg | Cloning Lin28a in pCI-H2B-GFP |
| 3´UTR-OC2-1_fwd.DL | taggcgatcgctcgaagcttgcgcagatgcacac | Cloning of 3´UTR-Dual luciferase |

| | | |
|---|---|---|
| 3´UTR-OC2-1_rev.DL | tagggggggggctcgatatgggttgtcaccaatgcg | Cloning of 3´UTR-Dual luciferase |
| 3´UTR-OC2-2_fwd.DL | taggcgatcgctcgagatccccggggcttctataaag | Cloning of 3´UTR-Dual luciferase |
| 3´UTR-OC2-2_rev.DL | tagggggggggctcgaaggggtggggctaggatg | Cloning of 3´UTR-Dual luciferase |
| 3´UTR-Pou2f2-1_fwd.DL | taggcgatcgctcgagatccccctcctcctcttcc | Cloning of 3´UTR-Dual luciferase |
| 3´UTR-Pou2f2-1_rev.DL | tagggggggggctcgagccttccacgctgcctgt | Cloning of 3´UTR-Dual luciferase |
| 3´UTR-Pou2f2-2_fwd.DL | taggcgatcgctcgagatccccctataggggagggg | Cloning of 3´UTR-Dual luciferase |
| 3´UTR-Pou2f2-2_rev.DL | tagggggggggctcgacagcccgagaaggcgggg | Cloning of 3´UTR-Dual luciferase |
| 3´UTR-Zfhx3_fwd.DL | taggcgatcgctcgagatccccctggccgcctc | Cloning of 3´UTR-Dual luciferase |
| 3´UTR-Zfhx3_rev.DL | tagggggggggctcgacgcacgggcttgcttctttg | Cloning of 3´UTR-Dual luciferase |
| Actin_F | TGGCTCCTAGCACCATGA | RT-qPCR |
| Actin_R | CCACCGATCCACACAGAG | RT-qPCR |

**Table S2.**

List of embryo genotypes used for scRNAseq.

| Sample | Breeding number | Embryo | Genotype | | Stage | Category |
|---|---|---|---|---|---|---|
| Brn4-Cre Dgcr8 flox | VP4700 | e1 | Brn4-Cre Dgcr8 hom | Cre + ; Dgcr8 flox/flox | e13.5 | mutant |
| Brn4-Cre Dgcr8 flox | VP4700 | e7 | Brn4-Cre Dgcr8 hom | Cre + ; Dgcr8 flox/flox | e13.5 | mutant |
| Brn4-Cre Dgcr8 flox | VP4700 | e4 | Brn4-Cre Dgcr8 het | Cre + ; Dgcr8 wt/flox | e13.5 | control |
| Brn4-Cre Dgcr8 flox | VP4700 | e6 | Brn4-Cre Dgcr8 het | Cre + ; Dgcr8 wt/flox | e13.5 | control |
| Brn4-Cre Dgcr8 flox | VP4923 | e2 | Brn4-Cre Dgcr8 het | Cre+; Dgcr8wt/flox; Rosa26-YFP$^{tg/wt}$ | e11.5 | control |
| Brn4-Cre Dgcr8 flox | VP4923 | e4 | Brn4-Cre Dgcr8 hom | Cre+; Dgcr8flox/flox; Rosa26-YFP$^{tg/wt}$ | e11.5 | mutant |
| Brn4-Cre Dgcr8 flox | VP4923 | e12 | Brn4-Cre Dgcr8 hom | Cre+; Dgcr8flox/flox; Rosa26-YFP$^{tg/wt}$ | e11.5 | mutant |
| Brn4-Cre Dgcr8 flox | VP5052 | e2 | Brn4-Cre Dgcr8 hom | Cre+; Dgcr8flox/flox; Rosa26-YFP$^{tg/wt}$ | e13.5 | mutant |
| Brn4-Cre Dgcr8 flox | VP5052 | e3 | Brn4-Cre Dgcr8 hom | Cre+; Dgcr8flox/flox; Rosa26-YFP$^{tg/wt}$ | e13.5 | mutant |
| Brn4-Cre Dgcr8 flox | VP5052 | e1 | Brn4-Cre Dgcr8 het | Cre+; Dgcr8wt/flox; Rosa26-YFP$^{tg/wt}$ | e13.5 | control |
| Brn4-Cre Dgcr8 flox | VP5553 | e2 | Brn4-Cre Dgcr8 het | Cre+; Dgcr8wt/flox; Rosa26-YFP$^{tg/wt}$ | e11.5 | control |
| Brn4-Cre Dgcr8 flox | VP5553 | e4 | Brn4-Cre Dgcr8 hom | Cre+; Dgcr8flox/flox; Rosa26-YFP$^{tg/wt}$ | e11.5 | mutant |
| Brn4-Cre Dgcr8 flox | VP5553 | e6 | Brn4-Cre Dgcr8 het | Cre+; Dgcr8wt/flox; Rosa26-YFP$^{tg/wt}$ | e11.5 | control |

**Table S3.**
List of resources and materials used for this research.

| Reagent type (species) or resource | Designation | Source or reference | Identifier | Additional information |
|---|---|---|---|---|
| Primary Antibody | Anti-cleaved caspase 3 (cC3) (Rabbit polyclonal) | Cell signaling | Cat #9661 RRID:AB_2341188 | IHC (1:200) |
| Primary Antibody | Anti-Cre-recombinase (Guinea pig polyclonal) | Synaptic Systems | Synaptic Systems Cat# 257 004 RRID:AB_2782969 | IHC (1:500) |
| Primary Antibody | Anti-Ctip2 (rat monoclonal) | Abcam | Abcam #ab18465 RRID:AB_2064130 | IHC (1:1000) |
| Primary Antibody | Anti-Dgcr8 (Rabbit Recombinant Monoclonal) | Abcam | Abcam #ab191875 RRID:AB_2892625 | IHC (1:1000) |
| Primary Antibody | Anti-En1 (polyclonal guinea pig) | Susan Brenner-Morton, Columbia University | CU1714, RRID:AB_3313271 | IHC 1:2500 |
| Primary Antibody | Anti-Foxp2 (Rabbit monoclonal) | Abcam | Abcam Cat# ab16046 RRID:AB_2107107 | IHC (1:2500) |
| Primary Antibody | Anti-Lhx1 (rabbit polyclonal) | Susan Brenner-Morton, Columbia University | CU453, RRID:AB_2827967 | IHC 1:20000 |
| Primary Antibody | Anti-Lmx1b (Guinea-pig polyclonal) | Thomas Müller (MDC, Berlin, Germany) | T. Müller, Max Delbruck Center for Molecular Medicine; Berlin; Germany Cat# Lmx1b RRID:AB_2314752 | IHC (1:10000) |
| Primary Antibody | Anti-Meis2 (mouse monoclonal) | Santa Cruz Biotechnology | Santa Cruz Biotechnology #sc-515470 RRID:AB_3076386 | IHC (1:1000) |
| Primary Antibody | Anti-NeuN (rabbit polyclonal) | Milipore | Milipore #ABN78 RRID:AB_10807945 | IHC (1:1000) |

| | | | | |
|---|---|---|---|---|
| Primary Antibody | Anti-Neurod2 (rabbit polyclonal) | Abcam | Abcam #ab104430 RRID:AB_10975628 | IHC (1:1000) |
| Primary Antibody | Anti-Nfib (rabbit monoclonal) | Abcam | Abcam #ab186738 RRID:AB_2782951 | IHC (1:200) |
| Primary Antibody | Anti-Nfix (rabbit ployclonal) | Atlas Antibodies | Atlas Antibodies #HPA007533 RRID:AB_2667855 | IHC (1:200) |
| Primary Antibody | Alexa Fluor® 647 Mouse Anti-Nkx2.2 | BD Biosciences | BD Biosciences #564729 RRID:AB_2738922 | IHC (1:50?) |
| Primary Antibody | Anti-OC2 (sheep polyclonal) | R&D Systems | R&D systems #AF6294 RRID:AB_10640365 | IHC (1:500) |
| Primary Antibody | Anti-Oct1 (Pou2f1) (rabbit Recombinant Monoclonal) | Abcam | Abcam #ab178869 RRID: N/A | IHC (1:1000) |
| Primary Antibody | Anti-Oct2 (Pou2f2) (monoclonal rat) | BD-Pharmingen | BD-Pharmigen #562837 RRID: N/A | IHC (1:500) |
| Primary Antibody | Anti-Oct6 (Pou3f1) (Rabbit Polyclonal) | Abcam | Abcam Cat# ab272925 RRID: N/A | IHC (1:500) |
| Primary Antibody | Anti-Olig3 (guinea pig polyclonal) | Thomas Müller (MDC, Berlin, Germany) | T. Müller, Max Delbruck Center for Molecular Medicine; Berlin; Germany | IHC (1:10000) |
| Primary Antibody | Anti-phosphohistone 3 (pHH3) (Rat monoclonal) | Novus Bio | Novus Bio #NB600-1168 RRID: N/A | IHC (1:1000) |
| Primary Antibody | Anti-Phox2a (Rabbit polyclonal) | Christo Goridis, Institut de Biologie du Développement de Marseille, France | Christo Goridis, Institut de Biologie du Développement de Marseille, France RRID: N/A | IHC (1:1000) |
| Primary Antibody | Anti-Pou2f2 (Oct-2) (Rabbit polyclonal) | Abcam | Cat# 178679, RRID: N/A | IHC (1:1000) |
| Primary Antibody | Anti-Pou3f1 (Oct6) (Mouse monoclonal) | EMD Millipore | MABN738, RRID: N/A | IHC (1:500) |

| | | | | |
|---|---|---|---|---|
| Primary Antibody | Anti-Sox10 (Guinea-pig polyclonal) | Michael Wegner (FAU-Erlangen-Nürnberg, Erlangen, Germany) | RRID: AB_2721917 | IHC (1:1000) |
| Primary Antibody | Anti-Sox2 (E-4) (mouse monoclonal) | Santa Cruz Biotechnology | Santa Cruz Biotechnology #sc-365823 RRID:AB_10842165 | IHC (1:500) |
| Primary Antibody | Anti-Sox2 (goat polyclonal) | R&D systems | R&D systems #AF2018 RRID:AB_355110 | IHC (1:500) |
| Primary Antibody | Anti-Sox9 (goat polyclonal) | R&D Systems | R&D systems #AF3075 RRID:AB_2194160 | IHC (1:500) |
| Primary Antibody | Anti-Zfhx3 (sheep plyclonal) | R&D Systems | R&D systems #ATBF1 RRID:AB_2218119 | IHC (1:1000) |
| Primary Antibody | Anti-Zfhx4 (rabbit polyclonal) | Atlas Antibodies | Atlas Antibodies #HPA023837 RRID:AB_1859013 | IHC (1:1000) |
| Primary Antibody | Calb1 (chicken polyclonal) | Novus Biologicals | Novus Biologicals #NBP2-50028 RRID: N/A | IHC (1:1000) |
| Secondary Antibody | Alexa Fluor® 488 Anti-Chicken (donkey polyclonal) | Jackson Immuno Research | Jackson Immuno Research #703-545-155 RRID: N/A | IHC (1:1000) |
| Secondary Antibody | Alexa Fluor® 488 Anti-Goat(donkey polyclonal) | Life Technologies (Invitrogen) | Life Technologies (Invitrogen) #A11055 RRID: N/A | IHC (1:1000) |
| Secondary Antibody | Alexa Fluor® 488 Anti-Guinea pig (donkey polyclonal) | Jackson Immuno Research | Jackson Immuno Research #706-545-148 RRID:AB_2340472 | IHC (1:1000) |
| Secondary Antibody | Alexa Fluor® 488 Anti-Mouse (donkey polyclonal) | Jackson Immuno Research | Jackson Immuno Research #706-545-148 RRID: N/A | IHC (1:1000) |

| | | | | |
|---|---|---|---|---|
| Secondary Antibody | Alexa Fluor® 488 Anti-Rabbit (donkey polyclonal) | Life Technologies (Invitrogen) | Life Technologies (Invitrogen) #A21206 | IHC (1:1000) |
| Secondary Antibody | Alexa Fluor® 488 Anti-Sheep (donkey polyclonal) | Life Technologies (Invitrogen) | Life Technologies (Invitrogen) #A11015 RRID:RRID:AB_141362 | IHC (1:1000) |
| Secondary Antibody | Alexa Fluor® 568 Anti-Goat(donkey polyclonal) | Life Technologies (Invitrogen) | Life Technologies (Invitrogen) #A11057 RRID:AB_142581 | IHC (1:1000) |
| Secondary Antibody | Alexa Fluor® 568 Anti-Mouse (donkey polyclonal) | Life Technologies (Invitrogen) | Life Technologies (Invitrogen) #A10037 RRID:AB_11180865 | IHC (1:1000) |
| Secondary Antibody | Alexa Fluor® 568 Anti-Rabbit (donkey polyclonal) | Invitrogen by Thermo Fisher Scientific | Invitrogen by Thermo Fisher Scientific #A10042 RRID: RRID:AB_2534017 | IHC (1:1000) |
| Secondary Antibody | Alexa Fluor® 568 Anti-Sheep (donkey polyclonal) | Life Technologies (Invitrogen) | Life Technologies (Invitrogen) #A21099 AB_2535753 | IHC (1:1000) |
| Secondary Antibody | Alexa Fluor® 647 Anti-Goat (donkey polyclonal) | Life Technologies (Invitrogen) | Life Technologies (Invitrogen) #A21447 RRID:AB_2535864 | IHC (1:500) |
| Secondary Antibody | Alexa Fluor® 647 Anti-Guinea pig (donkey polyclonal) | Jackson Immuno Research | Jackson Immuno Research #706-605-148 RRID:AB_2340476 | IHC (1:500) |
| Secondary Antibody | Alexa Fluor® 647 Anti-Mouse (donkey polyclonal) | Life Technologies (Invitrogen) | Life Technologies (Invitrogen) #A31571 RRID: N/A | IHC (1:500) |

| | | | | |
|---|---|---|---|---|
| Secondary Antibody | Alexa Fluor® 647 Anti-Rabbit (donkey polyclonal) | Life Technologies (Invitrogen) | Life Technologies (Invitrogen) #A31573 RRID:AB_2536183 | IHC (1:500) |
| Secondary Antibody | Alexa Fluor® 647 Anti-Rabbit (donkey polyclonal) | Invitrogen by Thermo Fisher Scientific | Invitrogen by Thermo Fisher Scientific #A21448 RRID: N/A | IHC (1:500) |
| Secondary Antibody | Alexa Fluor® 647 Anti-Rabbit (donkey polyclonal) | Invitrogen by Thermo Fisher Scientific | Invitrogen by Thermo Fisher Scientific #A78947 RRID: N/A | IHC (1:500) |
| Secondary Antibody | Alexa Fluor® 647Anti-Rabbit (donkey polyclonal) | Invitrogen by Thermo Fisher Scientific | Invitrogen by Thermo Fisher Scientific #A31573 RRID:AB_2536183 | IHC (1:500) |
| Secondary Antibody | Cy3-AffiniPure Donkey Anti-Rat IgG (H+L) | Jackson Immuno Research | Jackson Immuno Research # 712-165-153 RRID:AB_2340667 | IHC (1:200) |
| RNA scope probe | miRNAscope™ Probe - SR-mmu-let-7a-5p-S1 | Biotechne | Biotechne #727761-S1 RRID: N/A | |
| RNA scope probe | miRNAscope™ Probe - SR-mmu-miR-9-5p-S1 | Biotechne | Biotechne #1204801-S1 RRID: N/A | |
| RNA scope probe | RNAScope probe-Mm-Mir9-2 hg | Biotechne | Biotechne #1691171-C2 RRID: N/A | |
| RNA scope probe | RNAScope probe-Mm-Nfib-C2 | Biotechne | Biotechne #586511-C2 RRID: N/A | |
| RNA scope probe | RNAScope probe-Mm-pou2f2-O1-C2 | Biotechne | Biotechne #1170721-C2 RRID: N/A | |
| RNA scope probe | RNAScope probe-Mm-Zfhx3-O1-C3 | Biotechne | Biotechne #833461-C3 RRID: N/A | |
| RNA scope probe | RNAScope probe-Mm-Onecut2-C2 | Biotechne | Biotechne #520541-C2 RRID: N/A | |

| | | | |
|---|---|---|---|
| Plasmid | pCAG-nDsRedFL-intron | Addgene | Addgene #177804 RRID:Addgene_177804 |
| Plasmid | pCAG-nDsRedFL-miR9/9* | Addgene | Addgene #177805 RRID:Addgene_177805 |
| Plasmid | pCAG-nDsRedFL-miR9_sponge | Cloning | N/A |
| Plasmid | pCAG-nDsRedFL-let7 | Cloning | N/A |
| Plasmid | pCI-Lin28a-IRES-H2B-GFP | Cloning | N/A |
| Plasmid | psiCHECK3-Unstructured | Addgene | Addgene #136025 RRID:Addgene_136025 |
| Plasmid | psiCHECK3-3′UTR-OC2.1 | Cloning | N/A |
| Plasmid | psiCHECK3-3′UTR-OC2.2 | Cloning | N/A |
| Plasmid | psiCHECK3-3′UTR-Pou2f2.1 | Cloning | N/A |
| Plasmid | psiCHECK3-3′UTR-Pou2f2.2 | Cloning | N/A |
| Plasmid | psiCHECK3-3′UTR-Pou2f1 | Cloning | N/A |
| Plasmid | psiCHECK3-3′UTR-Zfhx3 | Cloning | N/A |
| TaqMan probe | mmu-mir-9-1, Mm04227702_pri | Thermo Fischer Scientific | Thermo Fischer Scientific # 4427012 |
| TaqMan probe | mmu-mir-9-2, Mm03306269_pri | Thermo Fischer Scientific | Thermo Fischer Scientific # 4427012 |
| TaqMan probe | mmu-mir-9-3, Mm03307250_pri | Thermo Fischer Scientific | Thermo Fischer Scientific # 4427012 |
| TaqMan probe | hsa-miR-9 (5p) | Thermo Fischer Scientific | Thermo Fischer Scientific #4427975 #000583 |
| TaqMan probe | hsa-miR-9* (3p) | Thermo Fischer Scientific | Thermo Fischer Scientific #4427975 #002231 |
| TaqMan probe | snoRNA 234-control miRNA | Thermo Fischer Scientific | Thermo Fischer Scientific #4427975 #001234 |

| | | | |
|---|---|---|---|
| TaqMan probe | hsa-let-7-a (5p) | Thermo Fischer Scientific | Thermo Fischer Scientific #4427975 #000377 |
| TaqMan probe | hsa-let-7-c | Thermo Fischer Scientific | Thermo Fischer Scientific #4427975 #000379 |
| Chemical | Alexa647 Click-iT EdU Imaging Kit | Thermo Fisher Scientific | Invitrogen #C10340 |
| Chemical | DAPI | Fisher Scientific | #D1306 |
| Chemical | Dual-Glo(R) luciferase assay system | Promega | Promage #E2920 |
| Chemical | Transfection agent: Lipofectamine 3000 | Thermo Fischer Scientific | Thermo Fischer Scientific #L3000015 |
| Chemical | Opti-MEM (100mL) | Thermo Fischer Scientific | Thermo Fischer Scientific # 31985062 |
| Chemical | TaqMan Fast advanced master mix | Applied Biosystems | Applied Biosystems #4444557 |
| Chemical | NEBuilder HiFi DNA Assembly Cloning Kit | New England Biolabs | New England Biolabs #E5520S |
| Chemical | PowerUp™ SYBR™ Green Master Mix | Thermo Fischer Scientific | Thermo Fischer Scientific #25743 |
| Chemical | cDNA synthesis kit with random primers - High capacity cDNA Reverse Transcription kit with Rnases Inhibitor | Thermo Fischer Scientific | Thermo Fischer Scientific #4374966 |
| Chemical | TaqMan microRNA Reverse Transcription kit | Thermo Fischer Scientific | Thermo Fischer Scientific #4366597 |
| Chemical | In-fusion Snap Assembly Master Mix | Takara | Takara #638948 |

**Table S4.**
List of the quantification methods and n number for each figure panel.

| Figure / Panel | Quantification | *N* | | Definition of ROIs |
|---|---|---|---|---|
| | | Embryo | Section/ embryo | |
| **1B** | *Dgcr8* cKO: number of positive nuclei (1n = 1 embryo) | WT = 6; cKO = 3 | 1-3 | Nuclei segmentation using StarDist + manual correction Thresholds: Pou2f2 = 0.175; Zfhx3 and Neurod2 = 0.2. Quantile = 0.001 |
| **1C** | *Dgcr8* cKO: fraction tTF double-positive/total tTF (1n = 1 embryo) | WT = 6; cKO = 3 | 1-3 | Nuclei segmentation using StarDist + manual correction |
| **1E** | *Dgcr8* cKO: number of positive nuclei (1n = 1 embryo) | WT = 6; cKO = 3 | 1-3 | Nuclei segmentation using StarDist + manual correction Thresholds: Pou2f2 = 0.175; OC2 = 0.25; Pou3f1 = 0.2. Quantile = 0.001 |
| | | Embryo | Section/ embryo | |
| **2B** | *Dgcr8* cKO: mean number of cells (1n = 1 embryo) | WT = 4; cKO = 4 | 2-3 | Nuclei segmentation using StarDist + manual correction |
| **2C** | *Dgcr8* cKO: fraction tTF+EdU+ double-positive/total EdU+ (1n = 1 embryo) | | | |
| OC2 / Pou2f2 | | WT = 4; cKO = 4 | 2-3 | Nuclei segmentation using StarDist + manual correction Thresholds: Pou2f2 = 0.175; OC2 = 0.25; EdU = 0.2. Quantile = 0.001 |
| Pou2f1 / Zfhx3 | | WT = 4; cKO = 4 | 2-3 | Nuclei segmentation using StarDist + manual correction Thresholds: Pou2f1 = 0.175; Zfhx3 = 0.2; EdU = 0.2. Quantile = 0.001 |
| **2E** | *Dgcr8* cKO: mean number of cells (1n = 1 embryo) | WT = 4; cKO = 4 | 2-3 | Nuclei segmentation using StarDist + manual correction |
| **2F** | *Dgcr8* cKO: fraction tTF+EdU+ double-positive/total Edu+ (1n = 1 embryo) | | | |
| OC2 / Pou2f2 | | WT = 6; cKO = 4 | 2-3 | Nuclei segmentation using StarDist + manual correction Thresholds: Pou2f2 = 0.175; OC2 = 0.25; EdU = 0.2. Quantile = 0.001 |

| | | | | |
|---|---|---|---|---|
| Pou2f1 / Zfhx3 | | WT = 6; cKO = 4 | 3 | Nuclei segmentation using StarDist + manual correction Thresholds: Pou2f1 = 0.175; Zfhx3 and EdU = 0.2. Quantile = 0.001 |
| **4H** | *Dgcr8* cKO: Normalised mean pixel intensity in Sox9+ cells (1n = 1 embryo) | Embryo | Section/ embryo | |
| Nfib/Sox10 | | WT = 6; cKO = 3 | 1 | Nuclei segmentation using StarDist + manual correction Quantile = 0.001 |
| Nfix | | WT = 6; cKO = 3 | 1 | Nuclei segmentation using StarDist + manual correction Quantile = 0.001 |
| Pou2f1 | | WT = 6; cKO = 3 | 1 | Nuclei segmentation using StarDist + manual correction Quantile = 0.001 |
| **7 B-D** | miR9 electroporation: Percentage of neurons positive for Pou2f1, Zfhx3, Bcl11b and total number of Neurod2 and mean intensity signal for Pou2f1, Zfhx3, Bcl11b and Neurod2 (1n = 1 embryo) | Embryo | Section/ embryo | |
| Bcl11b | | 5 | 2 | Nuclei segmentation using StarDist + manual correction Thresholds: Bcl11b = 0.125; Dsred = 0.1. Quantile = 0.01 for percentage and 0.05 for intensities |
| Pou2f1 | | 5 | 1-3 | Nuclei segmentation using StarDist + manual correction Thresholds: Pou2f1 = 0.125; Dsred = 0.1. Quantile = 0.01 for percentage and 0.05 for intensities |
| Zfhx3 | | 4 | 2-3 | Nuclei segmentation using StarDist + manual correction Thresholds: Zfhx3 = 0.125; Dsred = 0.1. Quantile = 0.01 for percentage and 0.05 for intensities |
| Neurod2 | | 4 | 2 | Cells counted manually using Fiji/ImageJ for total number & Nuclei segmentation using StarDist + manual correction for intensities Thresholds: Neurod2 = 0.125; Dsred = 0.1. Quantile = 0.05 for intensities |

| | | | | |
|---|---|---|---|---|
| **7 F, H** | miR9 sponge: Percentage of neurons positive for Pou2f1, Zfhx3, Bcl11b and mean intensity signal for Pou2f1, Zfhx3 and Bcl11b (1n = 1 embryo) | Embryo | Section/ embryo | |
| Bcl11b | | 3 | 2-5 | Nuclei segmentation using StarDist + manual correction Thresholds: Bcl11b = 0.125; Dsred = 0.1. Quantile = 0.05 |
| Pou2f1 | | 3 | 5-6 | Nuclei segmentation using StarDist + manual correction Thresholds: Pou2f1 = 0.125; Dsred = 0.1. Quantile = 0.05 |
| Zfhx3 | | 3 | 2-6 | Nuclei segmentation using StarDist + manual correction Thresholds: Zhfx3 = 0.125; Dsred = 0.1. Quantile = 0.05 |
| **7 G** | miR9 sponge: intensities of DsRed and Pou2f1 signal along the DV axis | Embryo | Section/ embryo | Nuclei segmentation using StarDist + manual correction |
| Pou2f1 | | 1 | 1 | Intensity profile measured in Fiji/ImageJ |
| | | Embryo | Section/ embryo | |
| **S2B** | *Dgcr8* cKO: Number of pHH3 positive cells (1n = 1 embryo) | WT = 3; cKO = 5 | 3-11 | Cells counted manually using Fiji/ImageJ for total number |
| **S2C** | *Dgcr8* cKO: Number of cC3 positive cells (1n = 1 embryo) | WT = 3; cKO = 5 | 3-11 | Cells counted manually using Fiji/ImageJ for total number |
| **S2D** | *Dgcr8* cKO: Percentage of areas: total, central canal, neuronal and progenitor (1n = 1 embryo) | WT = 3; cKO = 5 | 3-6 | Areas delineated manually using the Freehand selection tool in Fiji/ImageJ |
| **S2F** | *Dgcr8* cKO: Number of Sox2 and Sox2/Sox11 positive cells (1n = 1 embryo | WT = 3; cKO = 5 | 2-3 | Nuclei segmentation using StarDist + manual correction Thresholds: Sox2 and Sox11 = 0.2; Quantile = 0.001 |
| **S2H** | *Dgcr8* cKO: Percentage of areas: total, central canal, neuronal and progenitor (1n = 1 embryo) | WT = 6; cKO = 3 | 2-4 | Areas delineated manually using the Freehand selection tool in Fiji/ImageJ |
| **S2J** | *Dgcr8* cKO: Number of cC3 positive cells (1n = 1 embryo) | WT = 6; cKO = 3 | 4-11 | Cells counted manually using Fiji/ImageJ for total number |
| **S2K** | *Dgcr8* cKO: Number of pHH3 positive cells (1n = 1 embryo) | WT = 6; cKO = 3 | 4-11 | Cells counted manually using Fiji/ImageJ for total number |

| | | | | |
|---|---|---|---|---|
| **S4G** | *Dgcr8* cKO: mean number of cells / density (1n = 1 embryo) | WT = 6; cKO = 3 | 1-3 | Nuclei segmentation using StarDist + manual correction Thresholds: Phox2a and Zfhx3 = 0.2; Quantile = 0.001 |
| **S4I** | *Dgcr8* cKO: mean number of cells / density (1n = 1 embryo) | WT = 6; cKO = 3 | 1-3 | Nuclei segmentation using StarDist + manual correction Thresholds: Pou2f2 and Foxp1 = 0.2; Quantile = 0.001 |
| **S4K** | *Dgcr8* cKO: mean number of cells / density (1n = 1 embryo) | WT = 6; cKO = 3 | 1-3 | Nuclei segmentation using StarDist + manual correction Thresholds: Lhx1 and Olig3 = 0.2; Quantile = 0.001 |
| **S6** | Luciferase assay | | 4 biological replicate | Relative luminescence (Renilla / Firefly) and then normalized to avg of each |
| **S7** | RT-qPCR | | For miR-9 and pri-miR-9 5 biological replicates. For let-7 4 biological replicates | Delta delta CT values |
| **S9B** | let7 electroporation: Percentage of neurons positive for Pou2f1, Zfhx3, and Neurod2 (1n = 1 embryo) | Embryo | Section/embryo | |
| Neurod2 | | 5 | 2 | Cells counted manually using Fiji/ImageJ |
| Pou2f1 | | 4 | 3-5 | Nuclei segmentation using StarDist + manual correction Thresholds: Pou2f1 = 0.125; Dsred = 0.1. Quantile = 0.01 |
| Zfhx3 | | 3 | 2-3 | Nuclei segmentation using StarDist + manual correction Thresholds: Zfhx3 = 0.125; Dsred = 0.1. Quantile = 0.01 |
| **S9D** | Lin28a electroporation: Percentage of neurons positive for Pou2f1, Zfhx3, and Neurod2 (1n = 1 embryo) | Embryo | Section/ embryo | |
| Neurod2 | | 3 | 2-3 | Cells counted manually using Fiji/ImageJ |
| Pou2f1 | | 3 | 2-4 | Nuclei segmentation using StarDist + manual correction Thresholds: Pou2f1 = 0.125; Dsred = 0.1. Quantile = 0.01 |
| Zfhx3 | | 3 | 1-4 | Nuclei segmentation using StarDist + manual correction Thresholds: Zfhx3 = 0.125; Dsred = 0.1. Quantile = 0.01 |

| S9F | miR9 electroporation: Percentage of neurons positive for Meis2 and Zfhx4 (1n = 1 embryo) | Embryo | Section/ embryo | |
|---|---|---|---|---|
| Meis2 | | 4 | 2-3 | Nuclei segmentation using StarDist + manual correction Thresholds: Meis2 = 0.125; Dsred = 0.1. Quantile = 0.01 |
| Zfhx4 | | 4 | 2-3 | Nuclei segmentation using StarDist + manual correction Thresholds: Zfhx4 = 0.125; Dsred = 0.1. Quantile = 0.01 |
| **S9G** | intron electroporation: Percentage of neurons positive for Bcl11, Pou2f1, Zfhx3, Meis2 and Zfhx4 (1n = 1 embryo) | Embryo | Section/ embryo | |
| Bcl11b | | 5 | 2-4 | Nuclei segmentation using StarDist + manual correction Thresholds: Bcl11b = 0.125; Dsred = 0.1. Quantile = 0.01 |
| Pou2f1 | | 4 | 2-3 | Nuclei segmentation using StarDist + manual correction Thresholds: Pou2f1 = 0.125; Dsred = 0.1. Quantile = 0.01 |
| Zfhx3 | | 5 | 2-4 | Nuclei segmentation using StarDist + manual correction Thresholds: Zfhx3 = 0.125; Dsred = 0.1. Quantile = 0.01 |
| Meis2 | | 6 | 2-3 | Nuclei segmentation using StarDist + manual correction Thresholds: Meis2 = 0.125; Dsred = 0.1. Quantile = 0.01 |
| Zfhx4 | | 4 | 2-4 | Nuclei segmentation using StarDist + manual correction Thresholds: Zfhx4 = 0.125; Dsred = 0.1. Quantile = 0.01 |
